# GRAS salts and eastern hemlock extract: a dual approach to sustainable plant disease management

**DOI:** 10.1101/2025.11.07.687209

**Authors:** Veedaa Soltaniband, Adam Barrada, Maxime Delisle-Houde, Martine Dorais, Russell J. Tweddell, Dominique Michaud

## Abstract

Several studies assessed the potential of salts and plant extracts as sustainable, eco-friendly phytosanitary products in plant protection. Here, we examined the efficacy of Generally Recognized as Safe (GRAS) salts sodium benzoate (SBE) and sodium bicarbonate (SBI) used alone or in combination with a twig extract from eastern hemlock (EH) against the bacterial pathogen *Pseudomonas syringae* pv. *tomato* DC3000 (*Pst* DC3000). The antimicrobial activity of the three products, and their ability to bolster host plant natural defenses, were assessed using the Arabidopsis–*Pst* DC3000 pathosystem as a model. RT-qPCR and leaf staining assays on a transgenic Arabidopsis line engineered to express reporter protein ß-glucuronidase under the control of pathogen-inducible promoter *PR1* showed both GRAS salts and the EH extract to trigger the expression of salicylic acid (SA)-inducible genes. Salt- and EH extract-treated plants exhibited callose deposits in leaf tissue, further pointing to the induction of the SA signaling pathway and suggesting the establishment of a physical barrier in the leaf apoplast upon treatment. Direct toxicity measurements and a bacterial infection assay in growth chamber showed both GRAS salts and the EH extract to inhibit *Pst* DC3000 growth and Arabidopsis leaf colonization. Toxicity assays also revealed complementary, additive effects between the two salts and the EH extract in their detrimental effect against the pathogen, as evidenced by reduced minimum inhibitory concentrations when used in combination. Interestingly, SBE and SBI strongly delayed leaf senescence following infection, either alone or combined with the EH extract. Together, these findings confirm the potential of SBE, SBI and the EH twig extract as both toxic compounds against *Pst* DC3000 and natural defense stimulators in Arabidopsis. They also highlight the potential of GRAS salt and plant extract combinations as a sustainable alternative to chemical pesticides in plant pathogen management.

## 1 INTRODUCTION

Phytopathogenic bacteria affect a wide range of crops worldwide and cause diseases that lead to devastating damage and significant economic losses (Martins et al., 2018). The management of phytopathogenic bacteria in agriculture depends, in a large extent, on cultural practices and the heavy use of toxic chemicals. In Canada, the chemical control of pathogenic bacteria mainly relies on copper-based pesticides (Health Canada, 2025), which makes it imperative to develop low-risk substitutes given the reported negative effects of these products on human health, soil ecosystems and the environment (Schoffer et al., 2020; Mesquita et al., 2023). Further justifying the need for copper-based pesticide substitutes, numerous cases of resistance to copper have been reported over the years for different pathogenic bacteria, including from the widely distributed *Pseudomonas* (Alippi et al., 2003; Masami et al., 2004; Renick et al., 2008; Colombi et al., 2017), *Xanthomonas* (Vanneste et al., 2005; Behlau et al., 2012, 2013; Mirik et al., 2007) and *Erwinia* (Sholberg et al., 2001; Al-Daoude et al., 2009) genera. In this context, plant-derived extracts and ‘Generally Recognized as Safe’ (GRAS) products such as several salts used in the food industry have garnered interest in recent years for the development of low-risk substitutes to conventional pesticides (Bautista-Baños, 2014).

Several GRAS salts have shown efficiency to repress plant diseases. For instance, bicarbonate and carbonate salts were shown to suppress powdery mildew (*Sphaerotheca fuliginea*), gummy stem blight (*Didymella bryoniae*) and Alternaria leaf blight (*Alternaria cucumerina*) on cucurbit plants (Ziv & Zitter, 1992), powdery mildew (*Sphaerotheca pannosa* var. *rosae*) and black spot (*Diplocarpon rosae*) on roses (*Rosa* spp.) (Horst et al., 1992), green mold (*Penicillium digitatum*) on lemons (*Citrus limon*) and oranges (*Citrus sinensis*) (Smilanick et al., 1999), and potato tuber silver scurf (*Helminthosporium solani*) on potato (*Solanum tuberosum*) (Olivier et al., 1998; 1999; Hervieux et al., 2002). Potassium sorbate was reported to allow for the control of silver scurf on potato tubers (Olivier et al., 1998; 1999) and black root rot (*Thielaviopsis basicola*) on carrot (*Daucus carota*) (Punja & Gaye, 1993). Aluminum salts and sodium metabisulfite were shown to inhibit potato soft rot (*Pectobacterium atrosepticum*, *P. carotovorum*) (Yaganza et al., 2014), potato silver scurf (Hervieux et al., 2002) and potato dry rot (*Fusarium sambucinum*) (Mecteau et al., 2002), while sodium benzoate (SBE) was shown to repress potato soft rot (Yaganza et al., 2014) and Botrytis gray rot (*Botrytis cinerea*) on strawberry (*Fragaria* × *ananassa*) (El-Fawy et al., 2020). In these studies, a direct toxicity of the salts against the pathogens was confirmed but their impact on plant natural defense received little attention. Consequently, the efficacy of salts to control diseases was associated *de facto* with a direct toxic effect on the target pathogens, even though the stimulation of defense mechanisms by different salts had been demonstrated in some plants. For example, Jeandet et al. (2000) observed that aluminum chloride could stimulate the accumulation of resveratrol, a phytoalexin, in grapevine leaves, while callose synthesis was reported to be elicited by aluminum ions in several plants (Rengel, 1992).

Numerous studies have also reported the practical potential of plant extracts for plant pathogen control. Foliar sprays of garlic (*Allium sativum*), giant knotweed (*Reynoutria sachalinensis*) or licorice (*Glycyrrhiza glabra*) extracts on tomato (*Solanum lycopersicum*) plants were reported to repress early blight (*Alternaria solani*), powdery mildew (*Leveillula taurica*) and late blight (*Phytophthora infestans*), respectively (Konstantinidou-Doltsinis et al., 2006; Nashwa & Abo-Elyousr, 2012; Hermann et al., 2022). Likewise, garlic and neem (*Azadirachta indica*) extracts were reported to control potato late blight (*P. infestans*), while an extract from sodom (*Calotropis procera*) was shown to reduce the severity of potato tuber black scurf (*Rhizoctonia solani*) (Mahmood et al., 2022; Abdul-Karim & Hussein, 2025). Similarly, a sugar maple (*Acer saccharum*) autumn-shed leaf extract showed efficacy against lettuce (*Lactuca sativa*) varnish spot (*Pseudomonas cichorii*) and bacterial leaf spot (*Xanthomonas campestris* pv. *vitians*), angular leaf spot of cucurbits (*Pseudomonas syringae*) and tomato bacterial canker (*Clavibacter michiganensis* subsp. *michiganensis*) (Delisle-Houde & Tweddell, 2020; Tremblay et al., 2024; Mimouni et al., 2025). Higher plants contain a wide array of bioactive compounds, such as alkaloids, phenols, flavonoids and terpenes that show antimicrobial activity (Zuzarte et al., 2012; Mahizan et al., 2019; Tiku, 2020), and numerous studies have documented the detrimental effects of plant extracts against phytopathogenic fungi (Kishore et al., 2007; Nguefack et al., 2007; Bhagwat & Datar, 2014) and bacteria (Pradhanang et al., 2003; Ganiyu et al., 2017; Hermann et al., 2022).

Plant extracts are also increasingly recognized for their potency to stimulate defense mechanisms in plants (Aguilar-Gastélum et al., 2018; Shafique et al., 2019; Naz et al., 2021). For instance, seaweed extracts were shown to activate defense responses in a variety of plants including Arabidopsis (*Arabidopsis thaliana*) (Subramanian et al., 2011; Cook et al., 2018; Islam et al., 2020), strawberry (Hankins et al., 1990; Bajpai et al., 2019), tomato (Esserti et al., 2017; Ali et al., 2019; Melo et al., 2020), cucumber (*Cucumis sativus* var. *sativus*) (Jayaraman et al., 2011; Abkhoo & Sabbagh, 2016; Soliman et al., 2018), sweet pepper (*Capsicum annuum*) (Ali et al., 2019, 2021) and rice (*Oryza sativa*) (Flora & Rani, 2012), and to improve plant resistance against biotrophic and/or necrotrophic pathogens. Similarly, a giant knotweed (*Reynoutria sachalinensis*) extract was shown to reduce powdery mildew (*Podosphaera xanthii*) severity and spore germination in courgettes by activating defense responses including callose papillae formation, hydrogen peroxide synthesis and stimulation of the salicylic acid (SA)-dependent defense pathway (Margaritopoulou et al., 2020).

Recently, we the reported ethanolic extracts of forest trees, including eastern hemlock (EH) (*Tsuga canadensis*), English oak (*Quercus robur*), eastern red cedar (*Juniperus virginiana*) and red pine (*Pinus resinosa*), to both exhibit direct antibacterial effects against *P. syringae* pv. *tomato* strain *Pst* DC3000 and induce the expression of defense-related genes in leaf tissue of the model plant Arabidopsis (Soltaniband et al., 2024). Here, we investigated the plant protective potential of two GRAS salts, SBE and sodium bicarbonate (SBI), used alone or in combination with an EH extract, for their direct antibacterial activity and their ability to activate host plant natural defenses eventually detrimental to plant pathogens. We used Arabidopsis and *Pst* DC3000, the causal agent of tomato bacterial speck, as a model pathosystem (Xin & He, 2013).

## 2 MATERIALS AND METHODS

### 2.1 Plant materials and growth conditions

Arabidopsis transgenic line *PR1::GUS*, engineered to express reporter protein ß-glucuronidase (GUS) under the control of the SA-inducible promoter *PR1* (Koornneef et al., 2008), was used for the experiments. The seeds were surface sterilized, placed on a modified Murashige & Skoog medium (Soltaniband et al., 2024) and conditioned for 2 d at 4°C, before being transferred in a growth chamber for 14 d, until their roots reached the surface. The plants were kept under a light intensity of 80 μmol photons.m^−2^.s_−1_ during the day, a 16-h light photoperiod provided by fluorescent tubes, and a temperature regime of 23°C during the day and 18°C during the night. Water was provided as needed and replaced twice a week by a 50 ppm 20:20:20 NPK nutrient solution. For the bacterial infection assay (see below), 10 d-old seedlings were transferred in 6-cell trays containing Veranda Mix Potting Soil (Scotts Fafard, Saint-Bonaventure QC, Canada), and grown for 40 more days in the same conditions.

### 2.2 Chemicals

SBE (C_7_H_5_NaO_2_), sodium metabisulfite (Na_2_S_2_O_5_) and potassium sorbate (C_6_H_7_KO_2_) were purchased from Bio Basic Canada (Markham ON, Canada). Potassium bicarbonate (KHCO_3_) was from Ward’s Science (Rochester NY, U.S.A.), aluminum chloride (Al_2_Cl_3_) from Acros Organics (Geel, Belgium), SBI (NaHCO_3_) from Caledon (Georgetown ON, Canada), and sodium carbonate (Na_2_CO_3_) from Fisher Science Education (Rochester NY, U.S.A.). The SA functional mimic benzothiadiazole (BTH), commercialized as ACTIGARD^TM^ 50WG, was obtained from Syngenta Canada (Guelph ON, Canada).

### 2.3 Eastern hemlock extract

Twigs of eastern hemlock were collected on growing trees at Université Laval’s Roger-Van den Hende university garden (Québec QC, Canada). Ethanolic crude extracts were prepared as described by Delisle-Houde et al. (2020). The resulting powder was freeze-dried and stored in the dark at room temperature in Mason jars, before solubilization in sterile water for further use.

### 2.4 *Pst* DC3000

Rifampicin-resistant strain *Pst* DC3000 was used for the infection assay. The bacteria were maintained in 15% (v/v) glycerol (VWR International, West Chester PA, U.S.A.) until use, and then cultivated as needed at 28°C on King’s B (KB) solid medium, which included 20 g/L of Bacto™ Proteose Peptone No. 3 (Becton, Dickinson and Co., Sparks MD, U.S.A.) supplemented with 1.5 g/L dibasic sodium phosphate (EM Science, Gibbstown NJ, U.S.A.), 10 g/L glycerol (VWR International), 1.5 g/L magnesium sulfate (Fisher Scientific, Geel, Belgium) and 15 g/L CRITERION™ Agar (Hardy Diagnostics, Santa Maria CA, U.S.A.). The bacteria were inoculated in flasks containing 20 mL of KB liquid medium, incubated under agitation for 24 h at 28°C, recovered by centrifugation for 5 min at 3600×*g*, and resuspended in a 10 mM MgSO_4_ (Fisher Scientific) aqueous solution supplemented with 0.01% (w/v) SYLGARD™ OFX-0309 (Dow Chemical, Midland MI, U.S.A.), at a final concentration of 1×10^8^ cells/mL. Bacterial concentrations were monitored at 600 nm optical density (OD_600_) using 0.5 McFarland standards and an Epoch 2 Microplate Spectrophotometer (BioTek Instruments, Winooski VT, U.S.A.).

### 2.5 Activation of plant natural defenses

Gene induction assays were conducted to monitor defense gene expression in Arabidopsis seedlings treated with different salts (first assay) or with the most promising salts used alone or in combination with the EH extract (second assay). In the first assay, the salts were micro-filtrated through 0.22-μm filters and added at different doses (0.1 M, 0.01 M, and 0.001 M in 5 mL of water) to Petri dishes containing 14-d-old *PR1::GUS* Arabidopsis seedlings. In the second assay, the two most promising salts (SBE and SBI) were selected and used at a concentration of 0.01 M in 5 mL. The two salts were micro-filtered and added to Petri dishes containing 14-d old *PR1::GUS* Arabidopsis seedlings, either alone or in combination with the EH extract at a concentration of 25 mg/mL. For both assays, Petri dishes were incubated for 48 h at 22°C. Milli-Q water and ACTIGARD^TM^ 50WG at 0.5 mg/mL in sterile water were used as negative and positive controls, respectively. Following incubation, plant samples were taken for GUS activity assays and defense gene expression monitoring by quantitative reverse transcriptase (RT)-quantitative PCR (qPCR) using appropriate DNA primers (**Supplementary Table S1**). Three genes were selected for RT-qPCR, based on their differential inducibility by SA, JA and/or *Pst* DC3000 (Soltaniband et al., 2024) (**Table 1**).

**Table 1.** Defense marker genes for the qPCR assays.

| Gene marker | Locus <sup>1</sup> | Protein | Inducibility <sup>2</sup> |  |  |
| --- | --- | --- | --- | --- | --- |
|  |  |  | SA/BTH | JA/ET | DC3000 |
| *AtPR1* | at2g14610 | Pathogenesis-related protein 1 | • |  | • |
| *AtLOX2* | at3g45140 | Lipoxygenase 2 (13-LOX) |  | • |  |
| *AtPR3* | at3g12500 | Pathogenesis-related protein 3 |  | • | • |

<sup>1</sup> Accession numbers from The Arabidopsis Information Resource database (www.arabidopsis.org).
<sup>2</sup> As reported in The Arabidopsis Information Resource and/or the Arabidopsis GENEVESTIGATOR microarray database (Zimmermann et al., 2004).

### 2.6 GUS activity assay

Seedlings or plants at the blooming stage were used for the GUS activity assays. Leaf and seedling samples were transferred into microtubes containing 2 mL of GUS staining solution [1 mM X-Gluc (Biosynth International, San Diego CA, U.S.A.), 100 μM phosphate buffer (Fisher Scientific, Ottawa ON, Canada), 10 mM EDTA (Fisher Scientific), 0.1% (w/w) Triton (Fisher Scientific), 1 mM potassium ferrocyanide (II) (Sigma-Aldrich, Mississauga ON, Canada), 1 mM potassium ferricyanide (III) (Sigma-Aldrich)] and incubated for 16 h at 37°C. The samples were then washed in 70% (v/v) ethanol to remove the staining solution and then washed three more times for 1 h at 60°C in the same solvent to eliminate chlorophyll. Leaves and seedlings were observed under a stereomicroscope (Olympus Corporation Model SZ2-ILST, Tokyo, Japan). Images were captured and assembled using the photo merge tool of Adobe Photoshop CS6 (Adobe^®^ Photoshop). GUS staining intensities were quantified using the ImageJ software (Béziat et al., 2017).

### 2.7 RT-qPCR assays

RNA extraction and RT-qPCR were conducted as described in Soltaniband et al. (2024). Relative quantification of gene expression was performed using the PCR Miner software (Zhao & Fernald, 2005). Reference genes *Actin1 (ACT1)* and *Ubiquitin 5 (UBQ5*) were used for the normalization of gene expression. The stability of reference gene relative expression was validated according to Vandesompele et al. (2002), with an M threshold lower than 0.5 and a Cv threshold lower than 0.25.

### 2.8 Quantitation of callose deposition

Callose deposition was monitored in fresh seedling leaflets stained with aniline blue, following the procedure of Hael-Conrad et al. (2018). The leaflets were decolorized in a 96% (v/v) ethanol/diluted lactic acid mixture at a 3:1 volume ratio until all chlorophyll was extracted.

Translucent leaflets were rehydrated for 2 h in 50% (v/v) ethanol and then immersed for 1 h at 22°C in 67 mM K_2_HPO_4_, pH 12, containing 0.01% (w/v) aniline blue (Sigma). The stained samples were embedded in 30% (v/v) glycerol, placed on glass slides and examined under a Zeiss Axioskop 2 Plus (Zeiss Canada, Toronto ON, Canada) fluorescence microscope equipped with a blue excitation filter and a digital camera. Images of randomly selected fields were captured and processed with Adobe Photoshop CS6^®^ (Adobe, San Jose CA, U.S.A.) to estimate total areas of callose deposits. The procedure was performed in triplicate to ensure reliability and consistency.

### 2.9 Minimum inhibitory concentrations

Minimum inhibitory concentration (MIC) values were determined in sterile conditions using flat-bottom 96-well microplates (Sarstedt AG & Co., Nümbrecht, Germany), as described by Delisle-Houde et al. (2018). *Pst* DC3000 bacteria (5×10^5^ cells) were suspended in 100 μL of KB liquid nutrient medium containing different concentrations of salt (SBE or SBI) and EH extract. After incubation for 24 h at 28°C, 2,3,5-triphenyl-2H-tetrazolium chloride (Ward’s Science, Rochester NY, U.S.A.) was added to each well at a concentration of 1 mg/mL. MIC values corresponded to the lowest concentration of salt or plant extract (alone or in mixture) at which the bacteria showed no metabolic activity, i.e. no red coloration. The experiment was conducted twice, using three replicates for all salt and EH extract concentrations tested.

### 2.10 Fractional inhibitory concentration index values

Fractional inhibitory concentration (FIC) index values were calculated as described by Affia (2016) to identify eventual synergistic, additive or antagonistic interactions between the salts (SBE, SBI) and the EH extract in their antimicrobial effects against *Pst* DC3000. FIC index values were inferred from MIC values (as determined above) for each salt or EH extract treatment, used alone or in combination, using the following equation:

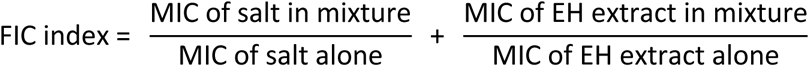

Calculated FIC index values were analyzed based on the scale of Zuo et al. (2014), where a FIC index lower than 0.5 indicates a synergistic interaction, a FIC index between 0.5 and 1.0 an additive interaction, a FIC index between 1.0 and 2.0 no interaction, and a FIC index greater than 2.0 an antagonistic interaction.

### 2.11 Bacterial infection assay

A bacterial infection assay was conducted in growth chamber using 30-day-old Arabidopsis *pPR1::GUS* plants at the blooming stage. Plants cultivated as described above were sprayed (20 mL per plant) with the EH extract (25 mg/mL), SBE (0.01 M), SBI (0.01 M), a combination of SBE and EH, a combination of SBI and EH, ACTIGARD™ 50WG (BTH; 0.5 mg/mL, positive control), or sterile water (negative control). The different formulations were applied two days prior to plant inoculation by dipping for 10 sec in a suspension of rifampicin-resistant *Pst* DC3000 mutant at 1 × 10⁸ cells/mL in 10 mM MgSO₄ containing 0.01% (w/v) SYLGARD™. Control (non-inoculated) plants were dipped in 10 mM MgSO₄ with 0.01% (w/v) SYLGARD™ only. Leaf samples were collected one day post *Pst* DC3000 inoculation for GUS activity detection, performed as described above. The plants were maintained in the growth chamber for an additional 10 days, after which they were harvested to assess leaf senescence, *Pst* DC3000 populations, and callose deposition. A leaf senescence index was determined for each plant using a scoring system based on the percentage of affected leaf area, where 0 = no visible symptom; 1 = more than 0% up to 25% affected leaf area; 2 = from 26% to 50% affected leaf area; 3 = from 51 to 75% affected leaf area; and 4 = more than 75% affected leaf area. The index values were calculated using the following formula:

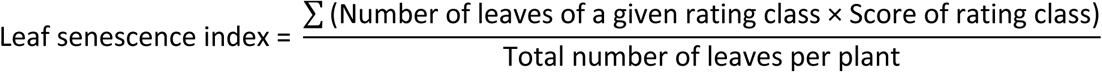

One week after inoculation, viable *Pst* DC3000 populations in leaves were quantified using a protocol adapted from Jacob et al. (2017). For each treatment, three leaves of similar size and age (totaling 200–300 mg) were randomly collected from three different plants within two replicates (two 6-cell seedling trays). The samples were homogenized twice for 30 seconds in 1 mL of 10 mM MgSO₄ using an Omni Bead Ruptor Homogenizer (Omni International Inc., Kennesaw, GA, USA). As described by Soltaniband et al. (2024), 10 µL of each bacterial suspension was serially diluted, and 10 µL of each dilution was spread onto KB agar plates supplemented with rifampicin at 50 mg/L (Sigma-Aldrich). Following incubation at 28 °C for 30 h, colony forming units (CFU) were counted, and *Pst* DC3000 populations expressed as CFU per milligram of fresh leaf tissue, relative to untreated control plants. The experiment followed a completely randomized design with three replicates, each consisting of one 6-cell seedling trays (six plants).

### 2.12 Statistical analyses

Analyses of variance (ANOVAs) or linear mixed-effects models (using the lme4 package) were performed using the R software, version 4.2.3 (https://www.r-project.org). Levene’s test (from the car package) and the Shapiro–Wilk test were used to assess homogeneity of variances and normality of the data, respectively. When necessary, raw data were transformed using Tukey’s ladder of powers (from the rcompanion package) or log transformation (*Pst* DC3000 populations) to improve conformity to normality assumptions. Treatment effects were compared using a post hoc LSD test (for ANOVAs) or pairwise comparisons of estimated marginal means with Tukey adjustment (from the emmeans package) for linear mixed-effects models.

## 3 RESULTS AND DISCUSSION

### 3.1 Selecting defense gene-inducing salts as a complement to the EH extract

Several studies have documented the direct toxic effects of GRAS salts against plant fungal and bacterial pathogens but less attention has been paid to the defense gene-inducing effects of these products *in planta*. An initial experiment was conducted with different salt formulations to identify salt candidates eventually effective in complementing the defense gene-inducing action of the EH extract (**Figure 1**). We recently reported a strong induction of GUS staining in seedlings of Arabidopsis *PR1::GUS* reporter line treated with this plant extract, attributable to an inducing effect on the transcriptional activity of the *PR1* promoter, an SA-inducible promoter driving the expression of pathogenesis-related protein PR1 upon microbial infection (Soltaniband et al., 2024). GRAS salts previously reported to show antimicrobial effects against plant pathogens, including SBE, SBI, aluminium chloride, sodium carbonate, potassium sorbate, sodium metabisulfite and potassium bicarbonate, were here tested with the same reporter line. As for the EH extract, all seven salts induced some GUS coloration of seedling tissues, albeit at varying intensities depending on the dose applied (**Figure 1A**).

**Figure 1.**
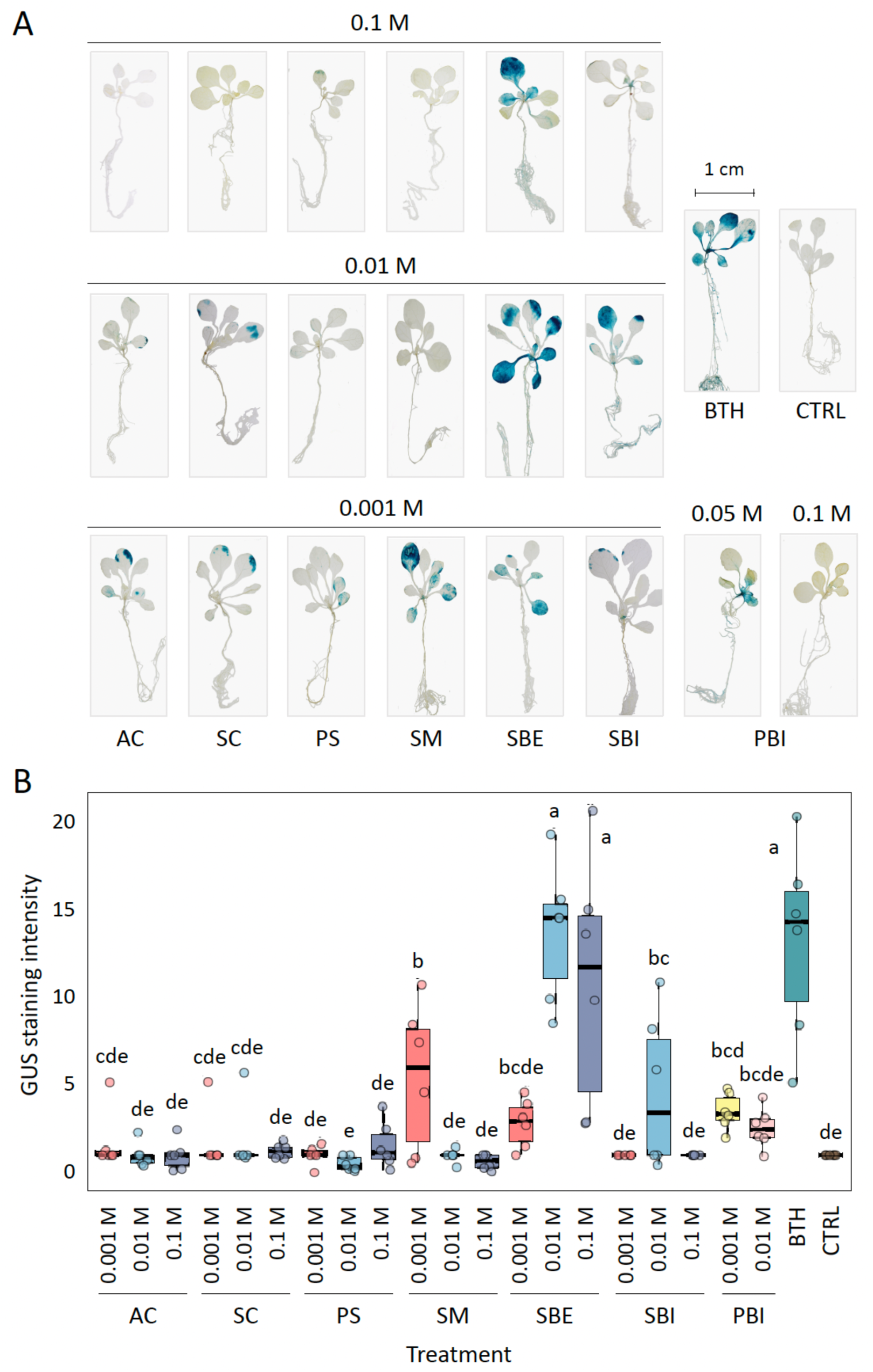
*PR1* promoter induction in seedlings of Arabidopsis reporter line *PR1::GUS* treated with different salt formulations. **A**, Representative images of salt-treated seedlings after GUS staining. **B**, Total GUS staining intensity per seedling, as estimated by image analysis with Adobe Photoshop™. The seedlings were treated with different concentrations of aluminum chloride (AC), sodium carbonate (SC), potassium sorbate (PS), sodium metabisulfite (SM), sodium benzoate (SBE), sodium bicarbonate (SBI) or potassium bicarbonate (PBI). Sterile water was used as a negative control treatment (CTRL), ACTIGARD^TM^ 50WG (BTH) at 0.5 mg/mL as a positive control for SA signaling induction. Individual measurements on panel B are shown as dots. In each box, the horizontal line shows the median, the lower and upper edges the first (Q1) and third (Q3) quartiles, respectively. Treatments sharing the same letter are not significantly different (n=6; *post-hoc* pairwise multiple comparisons of estimated marginal means with Tukey’s adjustment, with an alpha threshold value of 5%).

SBI (at 0.01 M), SBE (at 0.01 M and 0.1 M) and sodium metabisulfite (at 0.001 M) induced readily detectable GUS signals, stronger than the signals observed with the other salt treatments. Most noticeably, SBE induced GUS staining at levels comparable to SA mimic BTH (**Figure 1B**), consistent with the roles of benzoic acid as a trigger of SA-inducible responses and as a direct precursor of SA in many plants, including Arabidopsis (Ullah et al., 2023). Together, these observations indicating a clear induction of the *PR1* promoter in seedlings of SBE-, SBI- and sodium metabisulfite-treated plants suggested an eventual activating effect on the PR1 protein-encoding gene *in planta*. They also suggested the relevance of the three salts as a possible complement to plant extracts, including the EH extract, for the induction of host plant defenses. SBE, as a strong inducer of GUS staining in the reporter line, and SBI, as a moderate inducer of GUS staining, were selected for the next steps of the study.

### 3.2 SBE and SBI activate SA signaling in Arabidopsis seedlings

A second experiment was conducted with the *PR1::GUS* reporter line to compare the impacts of SBE, SBI and the EH extract on GUS staining, and to detect eventual interactions between these products in the treated seedlings (**Figure 2**). In accordance with GUS staining patterns above (**Figure 1A**) and the recently described inducing effects of the EH extract on *AtPR1* expression in Arabidopsis seedlings (Soltaniband et al., 2024), both the salts and the EH extract had a positive effect on GUS staining (**Figure 2A**), again as strong as for BTH in the SBE-treated seedlings (**Figure 2B**). By comparison, weaker GUS signals were observed for treatments combining SBE or SBI with the EH extract (*post-hoc* LSD test, *P* < 0.05), two to three times less intense than for BTH or SBE applied alone (**Figure 2B**). These observations suggested an interaction in treated seedlings between the salts and the plant extract, negatively influencing the net positive effect of either product on GUS production.

**Figure 2.**
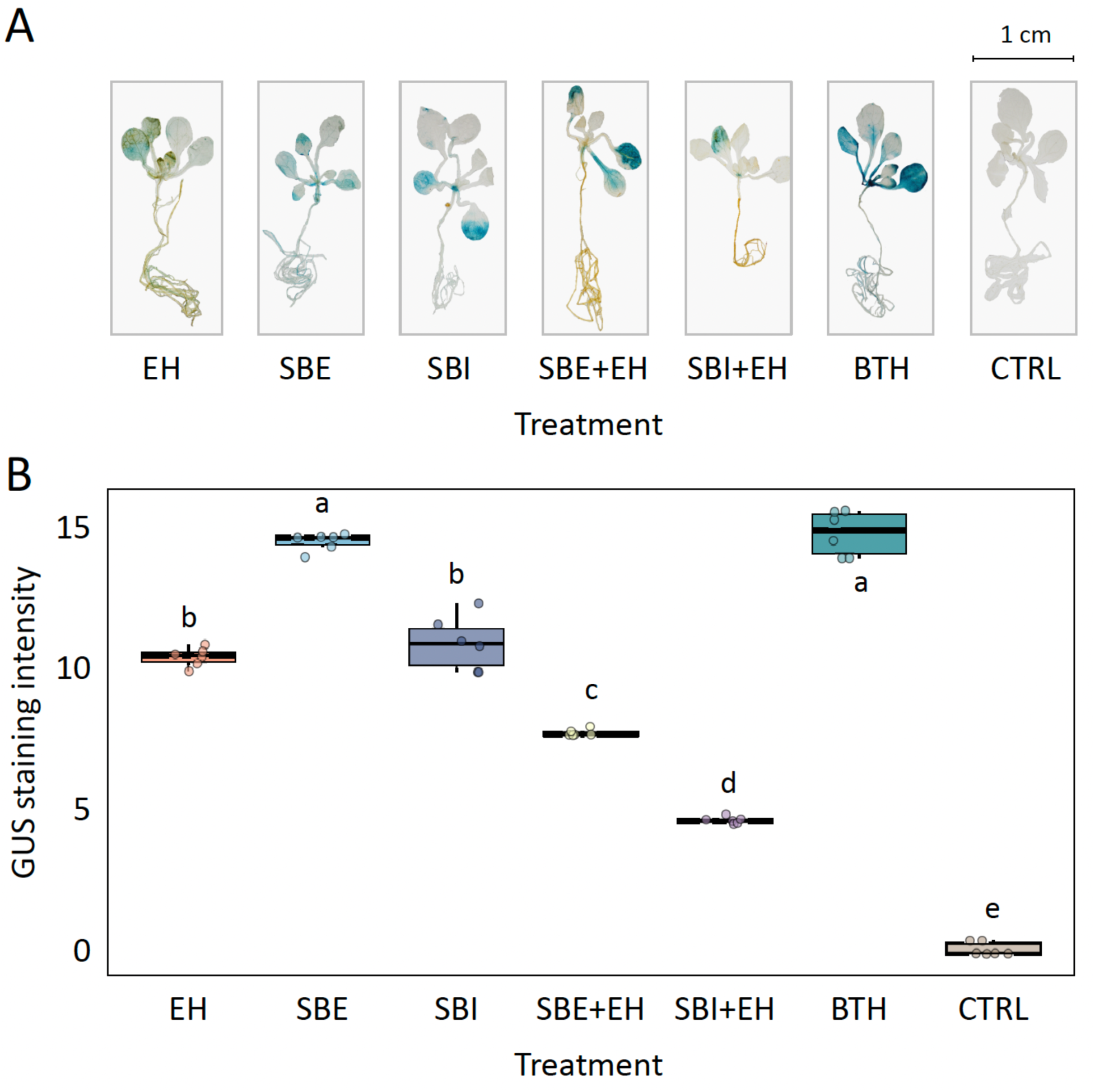
*PR1* promoter induction in seedlings of Arabidopsis reporter line *PR1::GUS* treated with sodium benzoate (SBE) or sodium bicarbonate (SBI), alone or combined with the eastern hemlock (EH) extract. **A**, Representative images of salt- and EH extract-treated seedlings after GUS staining. **B**, Total GUS staining intensity per seedling, as estimated by image analysis with Adobe Photoshop™. The seedlings were treated with 0.01 M SBE or 0.01 M SBI, with the EH extract at 25 mg/mL, or with SBE or SBI in combination with the EH extract at the same working concentrations. Sterile water was used as a negative control treatment (CTRL), ACTIGARD^TM^ 50WG (BTH) at 0.5 mg/mL as a non-toxic positive control for SA signaling induction. Individual measurements on panel B are shown as dots. In each box, the horizontal line shows the median, the lower and upper edges the first (Q1) and third (Q3) quartiles, respectively. Treatments sharing the same letter are not significantly different (n=6; *post-hoc* pairwise multiple comparisons of estimated marginal means with Tukey’s adjustment, with an alpha threshold value of 5%).

RT-qPCR assays were carried out to determine whether this interaction resulted from a reduced transcriptional activity of the *PR1* promoter or, alternatively, from a post- or co-translational effect affecting GUS deposition or stability in seedling tissues (**Table 2**). These assays were also performed to formally confirm the induction of SA signaling in salt-treated seedlings, and to look for a possible SA-independent induction of stress-related genes as recently suggested for the EH extract (Soltaniband et al., 2024). Three genes were used as markers, including SA-inducible gene *AtPR1* (encoding endogenous protein AtPR1), JA-inducible gene *AtLOX2* (encoding JA pathway lipoxygenase 2) and JA/ethylene-inducible *AtPR3* (encoding wound-inducible pathogenesis-related protein 3) (see **Table 1**). BTH, a non-toxic functional analogue of SA used as positive control for the SA signaling pathway (Görlach et al., 1996), strongly induced *AtPR1* transcription, as inferred from mRNA transcript numbers increased by almost 12-fold in BTH-treated seedlings compared to control seedlings (**Table 2**).

**Table 2.**
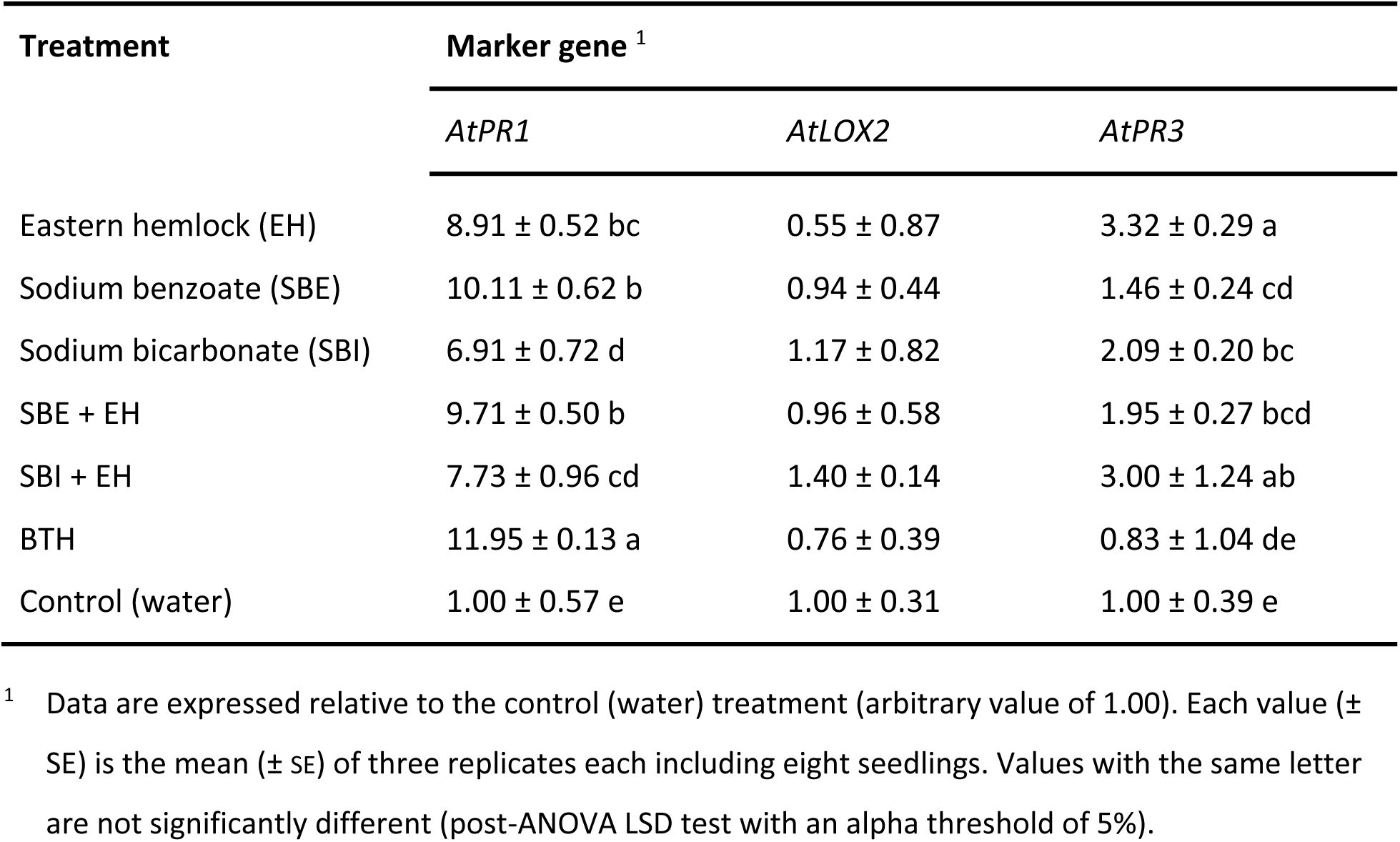
Expression of defense marker genes in Arabidopsis seedlings responding to eastern hemlock extract (EH) at a concentration of 25 mg/mL, to sodium benzoate (SBE) or sodium bicarbonate (SBI) at concentrations of 10 mM, and to combinations of SBE+EH or SBI+EH at the same concentrations. SA mimic ACTIGARD^TM^ 50WG (BTH) at 0.5 mg/mL was used as a positive control for SA-inducible gene *AtPR1*. Seedlings treated with sterile water were used as a negative control.

| Treatment | Marker gene <sup>1</sup> |  |  |
| --- | --- | --- | --- |
|  | *AtPR1* | *AtLOX2* | *AtPR3* |
| Eastern hemlock (EH) | 8.91 ± 0.52 bc | 0.55 ± 0.87 | 3.32 ± 0.29 a |
| Sodium benzoate (SBE) | 10.11 ± 0.62 b | 0.94 ± 0.44 | 1.46 ± 0.24 cd |
| Sodium bicarbonate (SBI) | 6.91 ± 0.72 d | 1.17 ± 0.82 | 2.09 ± 0.20 bc |
| SBE + EH | 9.71 ± 0.50 b | 0.96 ± 0.58 | 1.95 ± 0.27 bcd |
| SBI + EH | 7.73 ± 0.96 cd | 1.40 ± 0.14 | 3.00 ± 1.24 ab |
| BTH | 11.95 ± 0.13 a | 0.76 ± 0.39 | 0.83 ± 1.04 de |
| Control (water) | 1.00 ± 0.57 e | 1.00 ± 0.31 | 1.00 ± 0.39 e |

<sup>1</sup> Data are expressed relative to the control (water) treatment (arbitrary value of 1.00). Each value (± SE) is the mean (± SE) of three replicates each including eight seedlings. Values with the same letter are not significantly different (post-ANOVA LSD test with an alpha threshold of 5%).

As expected, given the GUS patterns above, *AtPR1* expression was also strongly induced by SBE, SBI and the EH extract, at relative levels reaching 7 to 10 times the expression levels measured in control plants. By comparison, *AtLOX2* was not induced by the salts or the EH extract, in sharp contrast with *AtPR3* showing transcript numbers increased by 46% to 232% in the same samples. For both *AtPR1* and *AtLOX2* but not for *AtPR3*, expression patterns for the salt and EH extract treatments matched the expression patterns observed with BTH. Unlike for GUS expression in the *PR1::GUS* reporter line (*see* **Figure 2B**), no negative effect was observed for the EH extract on the induction of *AtPR1* by SBE or SBI, suggesting the negative interaction inferred above between these elicitors to result from a post-transcriptional effect on GUS expression, rather than a downregulating effect on transcriptional activity of the *PR1* promoter.

We recently reported a strong activation of SA signaling in Arabidopsis seedlings treated with the EH extract, as inferred from the induction of SA-inducible genes *AtPR1*, transcription factor-encoding gene *AtWRKY70*, PR2 protein (ß-glucanase)-encoding gene *AtPR2* and Kunitz trypsin inhibitor-encoding gene *AtKTI4* (Soltaniband et al., 2024). Here we observed a significant inducing effect on *AtPR1* and a null effect on *AtLOX2* for SBE and SBI (**Table 2**), thus suggesting a similar SA signaling-promoting effect in treated seedlings. A likely explanation for this would be a generic, osmotic stress effect triggering SA signaling and *AtPR1* expression, as reported for other salts (Seo et al., 2008; Seo et al., 2020). An additional explanation in the case of SBE would be an inducing effect of its anion derivative, benzoic acid (C_6_H_5_COO^−^H^+^) (BA), on biosynthetic genes of the shikimate pathway, and the involvement of this metabolite in the plant as a direct precursor of SA along the phenyl ammonia lyase pathway (Lefevere et al., 2020; Peng et al., 2021; Ullah et al., 2023). Supporting this possibility, increased concentrations of SA and BA, along with a strong upregulation of three biosynthetic genes –isochorismate synthase, aldehyde oxidase and phenyl ammonia lyase– implicated in the production of these two elicitors were observed in tomato plants treated with BA or BA derivatives such as 4-hydroxybenzoic acid and protocatechuic acid (Nehela et al., 2021).

### 3.3 SBE and SBI also show an SA/JA-independent gene-inducing effect

Like for the EH extract (Soltaniband et al., 2024), an SA/JA-independent inducing effect on additional stress-related genes also remains likely for both SBE and SBI, given their upregulating effect on *AtPR3* expression and the null impact of BTH on this gene (**Table 2**). Dual induction of the SA and JA signaling pathways has been reported in Arabidopsis showing an effector-triggered immunity hypersensitive response (Betsuyaku et al., 2018) or treated with the polysaccharide elicitor chitosan (Jia et al., 2018), but this avenue remains unlikely in the present case given the divergent effects of JA and SA on *AtPR3* expression (Thomma et al., 1998; Seo et al., 2008) and the null impact of both salts on *AtLOX2* expression (**Table 2**). A more plausible avenue, considering the ionic and osmotic destabilizing effects of salts on cellular structures (Kesawat et al., 2023), would involve a yet to be elucidated, SA-independent oxidative stress response leading to the induction of *AtPR3* upon salt treatment (Seo et al., 2008).

Plants have evolved a complex array of antioxidative defenses to cope with salt stress, based on the production of non-enzymatic antioxidant compounds –such as phenolics and flavonoids– and reactive oxygen species (ROS)-scavenging enzymes –such as catalase, superoxide dismutase and ascorbate peroxidase– to protect themselves from oxidative damage (Zhou et al., 2024). In line with this, studies have reported increased levels of antioxidant compounds, higher activities of ROS-scavenging enzymes and/or increased numbers of transcripts for these enzymes in plants or plant-derived foods treated with various salts, including sodium silicate and potassium bicarbonate (Youssef et al., 2020), sodium carbonate (Ahmad et al., 2014; Youssef et al., 2014), sodium chloride (Hawrylak-Nowak et al., 2021), SBI (Youssef et al., 2014; Qin et al., 2017; Fan et al., 2021) and SBE (Moschetto et al., 2019; Rishad et al., 2021; Abo-Elyousr et al., 2022). Most interestingly, SBI was shown to trigger increased ß-glucanase (PR-2-like) and chitinase (PR-3-like) activities in potato leaves under field conditions (Abd-El-Kareem & Abd-El-latif, 2012), and sodium chloride to promote *AtPR3* transcription in an abscisic acid-dependent, SA-independent manner (Seo et al., 2008).

### 3.4 SBE, SBI and the EH extract trigger callose deposition in leaf tissue

Staining assays with aniline blue were conducted to detect an eventual accumulation of callose in SBE-, SBI- and EH extract-treated seedlings (**Figure 3**). Callose is a linear ß-(1-3)-D-glucan that plays various roles in plants, notably serving as a physical barrier to restrict the aperture of plasmodesmata and strengthening the cell wall during pathogenic infection (German et al., 2023). Leaf epidermal cells synthesize large amounts of callose upon infection, that affect the permeability of plasmodesmata and enhance resistance to several pathogens including *Pst* DC3000 (Liu et al., 2020). Callose deposition between the plasma membrane and the cell wall is strongly induced by SA in Arabidopsis, via an up-regulating effect on the biosynthesis of plasma membrane-located callose synthase isoforms catalyzing the addition of glucose units in the nascent polysaccharide (Wang et al., 2013). Consistent with Kohler et al. (2002) and as expected given the gene inducing effect of SA on callose synthase synthesis, a uniformly distributed aniline fluorescence pattern was observed in BTH-treated seedlings, much more intense than in the control treatment (**Figure 3A**,**B**).

**Figure 3.**
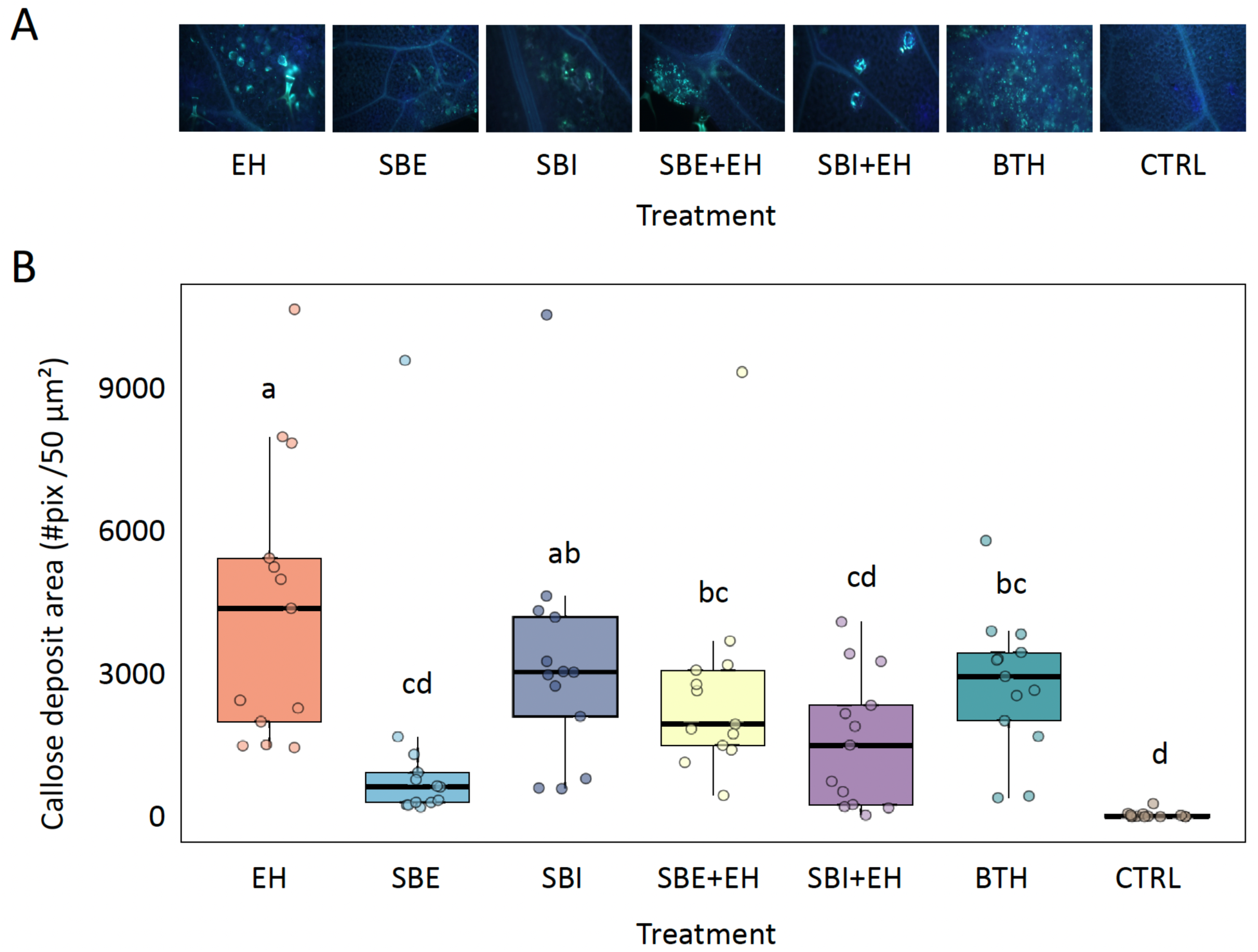
Callose deposition in seedling leaflets of Arabidopsis reporter line *PR1::GUS* treated with sodium benzoate (SBE) or sodium bicarbonate (SBI), alone or in combination with the eastern hemlock (EH) extract. **A**, Representative images of callose deposits following treatment, as visualized by aniline blue staining. **B**, Callose deposition intensity as quantified by the number of individual deposits per 50 µm^2^. The seedlings were treated with 0.01 M SBE or 0.01 M SBI, with the EH extract at 25 mg/mL, or with SBE or SBI combined with the EH extract at the same working concentrations. Sterile water was used as a negative control treatment (CTRL), ACTIGARD^TM^ 50WG (BTH) at 0.5 mg/mL as a non-toxic positive control for SA signaling induction. Individual measurements on panel B are shown as dots. In each box, the horizontal line shows the median, the lower and upper edges the first (Q1) and third (Q3) quartiles, respectively. Each mean value is based on measurements from 13 randomly selected seedling leaflet sections, derived from six plant replicates. Treatments sharing the same letter are not significantly different (*post-hoc* pairwise multiple comparisons of estimated marginal means with Tukey’s adjustment, with an alpha threshold value of 5%).

Similar to Llorens et al. (2019) who reported callose accumulation in lettuce plants upon treatment with other tree extracts, the EH extract strongly induced callose production in Arabidopsis seedlings, for an overall fluorescence intensity per unit area approximately 50% more intense than in BTH- and SBI-treated leaflets (**Figure 3B**). Unexpectedly, given the reported positive impact of SBE on SA and BE contents in leaves (Nehela et al., 2021), SBE-treated seedlings only showed a faint fluorescence signal, statistically comparable to the signal observed in seedlings assigned the control treatment and three to five times less than the fluorescence signals observed for BTH, SBI or the EH extract. As for the GUS signals above (**Figure 1**), salt treatments interfered with the staining-promoting effect of the EH extract, reducing aniline fluorescence signals by two- to three-fold compared to the EH extract alone (**Figure 3B**). Follow-up studies will be warranted to determine whether this interfering effect of SBE and SBI on callose induction by the EH extract was the result of a downregulating effect on callose synthase expression or, alternatively, of altered physicochemical conditions in the leaf cellular environment affecting cellulase synthase activity.

### 3.5 SBE, SBI and the EH extract are toxic to *Pst* DC3000 *in vitro*

Several studies have documented the toxic effects of SBE (e.g. El-Fawy et al., 2020; Williams et al., 2003; El-Mougy et al., 2004; Palou et al., 2009; Ragab et al., 2012; Montesinos-Herrero et al., 2016; Habibullah et al., 2020; Hussein & Ahmed, 2022) and SBI (e.g. Palmer et al., 1997; Smilanick et al., 1999; Sivakumar et al., 2002; Karabulut et al., 2003; De Costa & Gunawardhana, 2012; Fallanaj et al., 2016; Vilaplana et al., 2018; Lyousfi et al., 2022) against plant pathogens.

Based on this and considering the gene-inducing effects above, these two salts would potentially exert a dual protecting effect *in planta* encompassing direct toxic effects against the pathogen and indirect detrimental effects through their promoting impact on natural defenses in the host plant. To support this idea, we determined MIC and FIC index values to evaluate the direct toxicity of SBE (**Table 3**) and SBI (**Table 4**) against *Pst* DC3000, and to document potential interactions between the two salts and the EH extract administered in combination. MIC values, that here referred to the lowest concentration of active product (SBE, SBI or EH extract) at which no metabolic activity was observed for *Pst* DC3000 after an incubation of 24 h, were estimated at 1.73 mg/mL (0.012 M) and 4.2 mg/mL (0.050 M) for SBE and SBI, respectively, compared to 25 mg/mL for the EH extract (**Table 3**) and more than 50 mg/mL for BTH, a chemical not toxic to the pathogen (*see* Soltaniband et al., 2024). When used in combination, SBE and the EH extract exhibited reduced MIC values of 1.0 mg/mL (0.070 M) and 0.16 mg/mL, respectively (**Table 3**). Similarly, SBI and the EH extract used in combination showed reduced MIC values of 2.52 mg/mL (0.030 M mM) and 0.50 mg/mL, respectively (**Table 4**).

**Table 3.** Minimum inhibitory concentration (MIC) of sodium benzoate (SBE) and eastern hemlock (EH) extract, used alone or in combination, against *Pseudomonas syringae* pv. *tomato* DC3000, and calculated fractional inhibitory concentration (FIC) index.

| Treatment | MIC (mg/mL) <sup>1</sup> |
| --- | --- |
| SBE | 1.73 [0.012 M] |
| EH extract | 25 |
| SBE in combination | 1.00 [0.007 M] |
| EH extract in combination | 0.16 |
| FIC index (interaction) | 0.6 (additivity) |

<sup>1</sup> Each value is the mean of six replicate values. MIC of SBE in molarity reported in brackets.

**Table 4.**
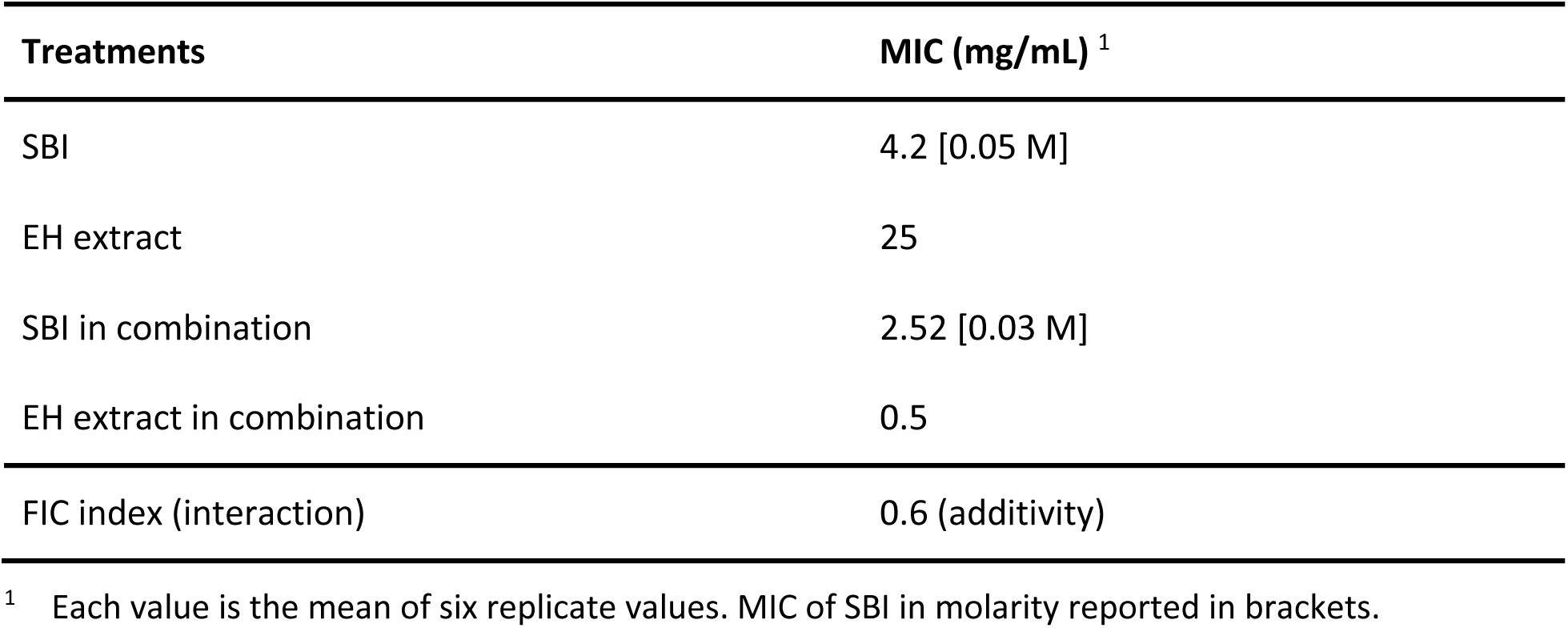
Minimum inhibitory concentration (MIC) of sodium bicarbonate (SBI) and eastern hemlock (EH) extract, used alone or in combination, against *Pseudomonas syringae* pv. *tomato* DC3000, and calculated fractional inhibitory concentration (FIC) index.

| Treatments | MIC (mg/mL) <sup>1</sup> |
| --- | --- |
| SBI | 4.2 [0.05 M] |
| EH extract | 25 |
| SBI in combination | 2.52 [0.03 M] |
| EH extract in combination | 0.5 |
| FIC index (interaction) | 0.6 (additivity) |

<sup>1</sup> Each value is the mean of six replicate values. MIC of SBI in molarity reported in brackets.

FIC index values were inferred from the MIC values to assess whether the salts and the EH extract showed interactions in their antibacterial effect when used in combination. FIC index values of 0.6 were determined for both the ‘SBE + EH extract’ (**Table 3**) and ‘SBI + EH extract’ combinations (**Table 4**), thereby pointing to additive effects between either salt and the plant extract in their toxicity against *Pst* DC3000. Previous studies have reported additive, or even synergistic, toxic effects between plant extracts and SBE used as a food or drug preservative against water- and foodborne pathogens such as *Staphylococcus aureus, Salmonella enterica* serovar Typhi*, Escherichia coli* or *Yersinia enterolitica* (Ekhtelat et al., 2020; Shaker et al., 2022), or against food spoiling bacteria like *Agrobacterium tumefaciens*, *Bacillus mycoides*, *Bacillus subtilis* or *Erwinia carotovora* (Stanojević et al., 2010). Here, we document additive toxic effects for SBE/SBI and the EH plant extract against *P. syringae*, likely explained by destabilizing effects on pH homeostasis, electrolyte balances, cell membrane integrity and metabolic functions once in the bacterium intracellular environment (Yaganza et al., 2009; Chipley, 2020).

### 3.6 SBE-, SBI- and EH extract-treated plants are resistant to *Pst* DC3000

Collectively, our data confirmed stress gene-inducing effects in Arabidopsis and direct toxic effects against *Pst* DC3000 for SBE and SBI, as recently reported for the EH extract (Soltaniband et al., 2024). An infection assay was carried out with the *PR1::GUS* reporter line to confirm the potential of SBE and SBI in preventing bacterial colonization of the host plant, and to evaluate the feasibility of using these salts in combination with the EH extract for *Pst* DC3000 control.

Leaf senescence index values were first determined to rule out the possibility of phytotoxic effects and to detect possible anti-senescence effects for the salt treatments following infection (**Figure 4**). In accordance with our recent work comparing the protective effects of BTH and different forest tree extracts on Arabidopsis against *Pst* DC3000 (Soltaniband et al., 2024), senescence index values for BTH- and EH extract-treated plants were roughly similar to the senescence index values of control plants, for an average index value of approximately 1.0 (**Figure 4B**) and a dozen leaves per plant presenting visible symptoms of necrosis (**Figure 4A**). By comparison, senescence index values close to 0.0, and plants presenting no visible symptom of necrosis, were observed after nine days on plants treated with SBE and SBI applied alone or in combination with the EH extract (**Figure 4B**). Similar observations were made with non-infected plants submitted to the same treatments, thus suggesting an anti-senescence effect for the two salts independent of *Pst* DC3000 infection (**Supplementary Figure S1**).

**Figure 4.**
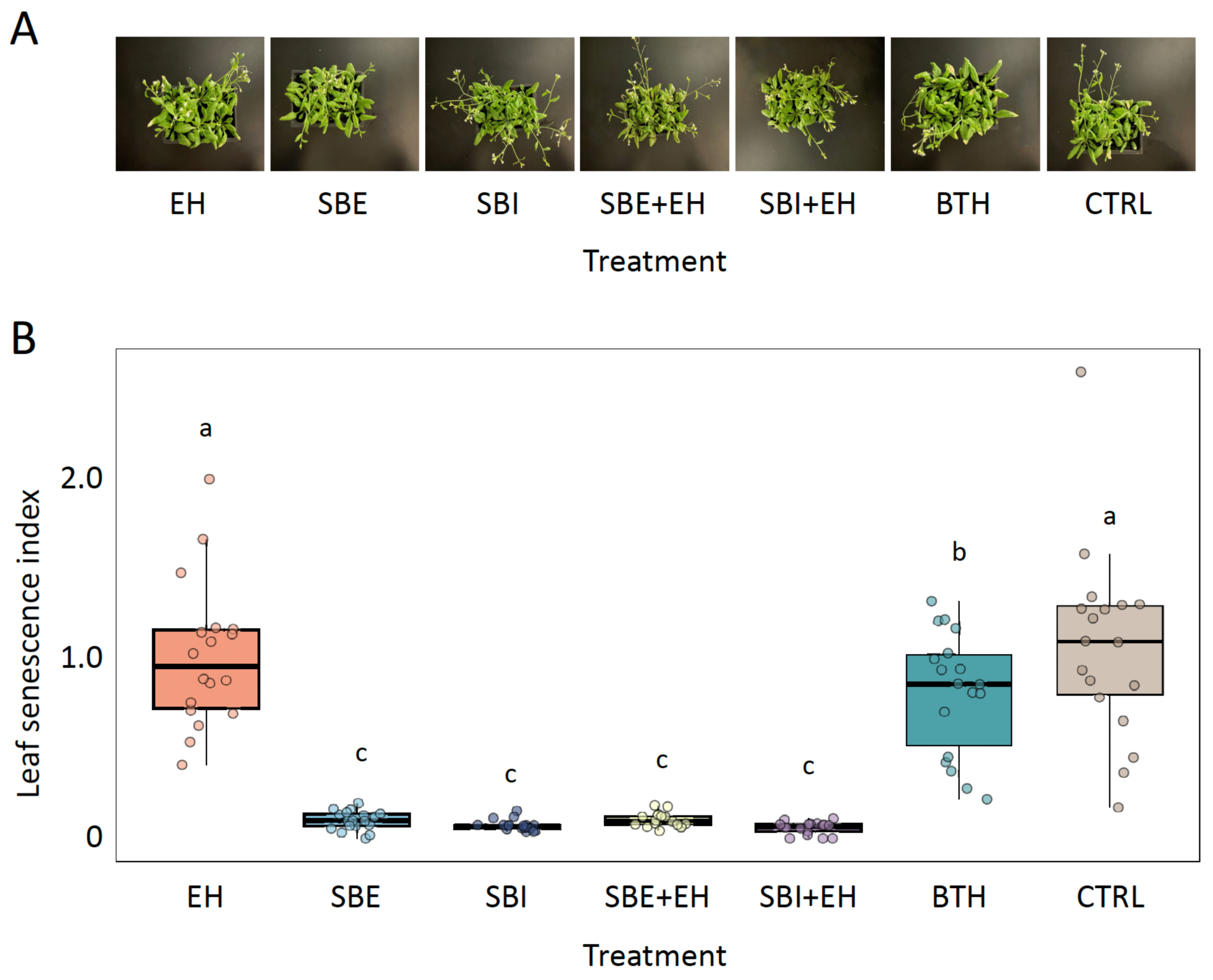
Leaf senescence index of Arabidopsis plants treated with sodium benzoate (SBE) or sodium bicarbonate (SBI), alone or combined with the eastern hemlock (EH) extract. **A**, Representative images of leaves displaying varying degrees of senescence. **B**, Leaf senescence levels, as quantified using the leaf senescence index. The plants were treated with 0.01 M SBE or 0.01 M SBI, with the EH extract at 25 mg/mL, or with SBE or SBI in combination with the EH extract at the same working concentrations. Sterile water was used as a negative control treatment (CTRL), ACTIGARD^TM^ 50WG (BTH) at 0.5 mg/mL as a non-toxic positive control for SA signaling induction. Treatments were applied by foliar spraying two days prior to inoculation with *Pst* DC3000, and the leaf samples collected after nine days. Individual measurements on panel B are shown as dots. In each box, the horizontal line shows the median, the lower and upper edges the first (Q1) and third (Q3) quartiles, respectively. Treatments sharing the same letter are not significantly different (n=18; *post-hoc* pairwise multiple comparisons of estimated marginal means with Tukey’s adjustment, with an alpha threshold value of 5%).

The plants were assessed for their resistance against *Pst* DC3000 to detect eventual differential antibacterial efficiencies for the different treatments, given the variable toxicity index values and gene-inducing effects described above for BTH, SBE, SBI and the EH extract. GUS staining assays conducted 7 days post-infection confirmed induction of the *PR1* promoter in salt- and EH extract-treated leaves, indicative of SA signaling induction by these treatments, albeit at levels lower than the BTH treatment (**Supplementary Figure S2**). *Pst* DC3000 populations were quantified in the same plant samples to assess the antibacterial potential of the salt and EH extract treatments, compared to BTH used as a non-toxic defense elicitor (**Table 5**). Despite variable GUS deposition rates in leaves, bacterial counts recovered from plants sprayed with SBE and SBI used alone or in combination with the EH extract were comparable to those observed for the BTH treatment, less than 6% the CFU counts observed for the control plants. Together, these observations indicated a negligeable induction of SA signaling by the bacterial pathogen under our conditions, and a GUS staining rate in BTH-treated leaves greater than in leaves assigned the salt treatments. Considering the strong antibacterial effects of the salt treatments despite weaker GUS signals, our observations also suggested a contribution of both the salt and EH extract direct toxic effects to the net protective effect of these products *in planta*.

**Table 5.** *Pst* DC3000 populations in leaf tissue of Arabidopsis plants treated with the eastern hemlock extract (EH), sodium benzoate (SBE), sodium bicarbonate (SBI), or a combination of EH extract and SBE (or SBI). ACTIGARD™ 50WG (BTH) and sterile water treatments were used as positive and negative controls, respectively. The plants were treated by foliar spraying two days prior to inoculation with *Pst* DC3000.

| Treatment | *Pst* DC3000 population (%) <sup>1</sup> |
| --- | --- |
| Eastern hemlock (EH) | 3.14 ± 1.74 ab |
| Sodium benzoate (SBE) | 3.27 ± 1.25 ab |
| Sodium bicarbonate (SBI) | 1.03 ± 0.09 b |
| SBE + EH | 5.75 ± 2.39 a |
| SBI + EH | 2.20 ± 0.07 ab |
| BTH | 2.36 ± 1.22 ab |

<sup>1</sup> Data are expressed as a percentage relative to water-treated plants (100%). Each value (± SE) is based on measurements from three randomly selected plants in two replicates. Means sharing the same letter are not significantly different (n = 6; pairwise comparison of estimated marginal means with Tukey's adjustment, with an alpha threshold value of 5%).

## 4 CONCLUSION

Several studies have discussed the antimicrobial potential of GRAS salts, including SBE and SBI, as a sustainable alternative to conventional pesticides in plant protection. Here, we assessed the potential of SBE and SBI against a well-characterized pathovar of *P. syringae*, a bacterial pathogen of worldwide economic importance in agriculture (Xin et al., 2018). Our findings highlighted a potential dual effect for the two salts, involving direct toxic effects against the pathogen and gene-inducing effects triggering at least two metabolic pathways of the host plant defense machinery. Our data also confirmed the feasibility of using SBE and SBI in combination with the EH extract, an ethanolic extract from twigs of eastern hemlock also identified as a possible eco-friendly alternative to conventional pesticides for *Pst* DC3000 management (Soltaniband et al., 2024). The two salts and the EH extract, while showing similar defense gene-inducing effects *in planta*, showed functional complementarity in terms of antisenescence potential and direct toxicity against the pathogen. A strong repression of bacterial colonization was observed for all salt/EH extract combinations one week after *Pst* DC3000 inoculation, comparable to the protective effect of BTH used as a non-toxic elicitor of SA-mediated responses. Complementary studies will be warranted in coming years to evaluate the relative contributions of direct toxicity and plant gene induction to the plant protecting effects of the two salts and the EH extract, and to confirm a possible dual effect for these products in bacteria-infected leaves. Studies will also be welcome to confirm the protective potential and complementary action of these products against other plant pathogens, including additional *P. syringae* pathovars of economic relevance (Chen et al., 2022).

## ACKNOWLEDGEMENTS

The Arabidopsis *PR1::GUS* line used in this study was generously provided by Prof. Corné Pieterse, Utrecht University, The Netherlands. This work was supported by a grant from the Ministère de l’agriculture, des pêcheries et de l’alimentation du Québec (Programme Innov’Action Agroalimentaire), with the involvement of Investissement Québec-CRIQ and Les Fraises de l’île d’Orléans inc.

## DATA AVAILABILITY STATEMENT

Data supporting the findings of this study are available from the corresponding authors upon reasonable request.

## SUPPORTING INFORMATION

Additional supporting information can be found online.

## REFERENCES

Abd-El-Kareem, F. & Abdul-El-latif, F.M. (2012) Using bicarbonates for controlling late blight disease of potato plants under field conditions. Life Science Journal, 9, 2080–2085.

Abdul-Karim, E. & Hussein, H.Z. (2025) Efficiency of aqueous and alcoholic extract of *Calotropis procera* in resisting the fungus *Rhizoctonia solani*, the causative agent of black scurf disease on potatoes. Polish Journal of Environmental Studies, 34, 2513–2523.

Abkhoo, J. & Sabbagh, S.K. (2016) Control of *Phytophthora melonis* damping-off, induction of defense responses, and gene expression of cucumber treated with commercial extract from *Ascophyllum nodosum*. Journal of Applied Phycology, 28, 1333–1342.

Abo-Elyousr, K.A.M., Imran, M., Almasoudi, N.M., Ali, E.F., Hassan, S., Sallam, N.M.A. et al. (2022) Controlling of *Xanthomonas axonopodis* pv. *phaseoli* by induction of phenolic compounds in bean plants using salicylic and benzoic acids. Journal of Plant Pathology, 104, 947–957.

Affia, H. (2016) Évaluation de différents sels et mélanges de sels pour lutter contre Pseudomonas cichorii dans la laitue. Master’s theses, Québec, Université Laval.

Aguilar-Gastélum, I., Martínez-Téllez, M.Á., Corrales-Maldonado, C., Rivera-Domínguez, M., Vargas-Arispuro, I. & Arellano-Gil, M. (2018) Induction of defense response in tomato plants against Forl by garlic extract. Revista Mexicana de Fitopatología, 36, 394–413.

Ahmad, P., Ozturk, M., Sharma, S. & Gucel, S. (2014) Effect of sodium carbonate-induced salinity–alkanility on some key osmoprotectants, protein profile, antioxidant enzymes, and lipid peroxidation in two mulberry (*Morus alba* L.) cultivars. Journal of Plant Interactions, 9, 460–467.

Al-Daoude, A., Arabi, M.I.E. & Ammouneh, H. (2009) Studying *Erwinia amylovora* isolates from Syria for copper resistance and streptomycin sensitivity. Journal of Plant Pathology, 91, 203– 205.

Ali, O., Ramsubhag, A. & Jayaraman, J. (2019) Biostimulatory activities of *Ascophyllum nodosum* extract in tomato and sweet pepper crops in a tropical environment. PLoS ONE, 14, e0216710.

Ali, O., Ramsubhag, A. & Jayaraman, J. (2021) Phytoelicitor activity of *Sargassum vulgare* and *Acanthophora spicifera* extracts and their prospects for use in vegetable crops for sustainable crop production. Journal of Applied Phycology, 33, 639–651.

Alippi, A.M., Dal Bo, E., Ronco, L.B., López, M.V., López, A.C. & Aguilar, O.M. (2003) *Pseudomonas* populations causing pith necrosis of tomato and pepper in Argentina are highly diverse. Plant Pathology, 52, 287–302.

Bajpai, S., Shukla, P.S., Asiedu, S., Pruski, K. & Prithiviraj, B. (2019) A biostimulant preparation of brown seaweed *Ascophyllum nodosum* suppresses powdery mildew of strawberry. The Plant Pathology Journal, 35, 406–416.

Bautista-Baños, S. (2014) Postharvest decay: Control strategies. Waltham, MA: Academic Press.

Behlau, F., Canteros, B.I., Jones, J.B. & Graham, H.J. (2012) Copper resistance genes from different xanthomonads and citrus epiphytic bacteria confer resistance to *Xanthomonas citri* subsp. *citri*. European Journal of Plant Pathology, 133, 949–963.

Behlau, F., Hong, J.C., Jones, J.B. & Graham, J.H. (2013) Evidence for acquisition of copper resistance genes from different sources in citrus-associated xanthomonads. Phytopathology, 103, 409–418.

Betsuyaku, S., Katou, S., Takebayashi, Y., Sakakibara, H., Nomura, N. & Fukuda, H. (2018) Salicylic acid and jasmonic acid pathways are activated in spatially different domains around the infection site during effector-triggered immunity in *Arabidopsis thaliana*. Plant and Cell Physiology, 59, 8–16.

Béziat, C., Kleine-Vehn, J. & Feraru, E. (2017) Histochemical staining of β-glucuronidase and its spatial quantification. In: Kleine-Vehn, J. & Sauer, M. (Eds.) Plant Hormones: Methods and Protocols. New York, NY: Humana Press, pp.73–80.

Bhagwat, M.K. & Datar, A.G. (2014) Antifungal activity of herbal extracts against plant pathogenic fungi. Archives of Phytopathology and Plant Protection, 47, 959–965.

Chen, H., Chen, J., Zhao, Y., Liu, F. & Fu, Z.Q. (2022) Pseudomonas syringae pathovars. Trends in Microbiology, 30, 912–913.

Chipley, J.R. (2020) Sodium benzoate and benzoic acid. In: Davidson, P.M., Taylor, T.M. & David, J.R.D. (Eds.) Antimicrobials in Food 4th edition. Boca Raton, FL: CRC Press, pp. 41–87.

Colombi, E., Straub, C., Künzel, S., Templeton, M.D., McCann, H.C. & Rainey, P.B. (2017) Evolution of copper resistance in the kiwifruit pathogen *Pseudomonas syringae* pv. *acbnidiae* through acquisition of integrative conjugative elements and plasmids. Environmental Microbiology, 19, 819–832.

Cook, J., Zhang, J., Norrie, J., Blal, B. & Cheng, Z. (2018) Seaweed extract (Stella Maris®) activates innate immune responses in *Arabidopsis thaliana* and protects host against bacterial pathogens. Marine Drugs, 16, 221.

De Costa, D.M. & Gunawardhana, H.M.D.M. (2012) Effects of sodium bicarbonate on pathogenicity of *Colletotrichum musae* and potential for controlling postharvest diseases of banana. Postharvest Biology and Technology, 68, 54–63.

de Melo, P.C., Collela, C.F., Sousa, T., Pacheco, D., Cotas, J., Gonçalves, A.M.M. et al. (2020) Seaweed-based products and mushroom β-glucan as tomato plant immunological inducers. Vaccines, 8, 524.

Delisle-Houde, M., Toussaint, V., Affia, H. & Tweddell, R.J. (2018) Evaluation of different salts for the control of lettuce varnish spot: when phytotoxicity rules. Canadian Journal of Plant Science, 98, 753–761.

Delisle-Houde, M., Dubé, P. & Tweddell, R.J. (2020) Antibacterial activity of sugar maple autumn-shed leaf extract: Identification of the active compound. Annals of Applied Biology, 177, 51–60.

Delisle-Houde, M. & Tweddell, R.J. (2020) Sugar maple autumn-shed leaf extract: a potential antibacterial agent for the control of lettuce bacterial leaf spot and varnish spot. Canadian Journal of Plant Science, 100, 78–85.

Ekhtelat, M., Borujeni, F.k., Siahpoosh, A. & Ameri, A. (2020) Chemical composition and antibacterial effects of some essential oils individually and in combination with sodium benzoate against methicillin-resistant *Staphylococcus aureus* and *Yersinia enterocolitica*. Veterinary Research Forum, 11, 333–338.

El-Fawy, M.M., El-Sharkawy, R.M.I. & Ahmed, M.M.Sh. (2020) Impact of pre- and post-harvest treatment with chemicals preservatives on Botrytis gray rot disease and fruit quality of strawberry. Archives of Agriculture Sciences Journal, 3, 178–194.

El-Mougy, N.S., Abd-El-kareem, F.A., El-Gamal, N.G. & Fotouh, Y.O. (2004) Application of fungicides alternatives for controlling cowpea root rot diseases under greenhouse and field conditions. Egyptian Journal of Phytopathology, 32, 23–35.

Esserti, S., Smaili, A., Rifai, L.A., Koussa, T., Makroum, K., Belfaiza, M. et al. (2017) Protective effect of three brown seaweed extracts against fungal and bacterial diseases of tomato. Journal of Applied Phycology, 29, 1081–1093.

Fallanaj, F., Ippolito, A., Ligorio, A., Garganese, F., Zavanella, C. & Sanzani, S.M. (2016) Electrolyzed sodium bicarbonate inhibits *Penicillium digitatum* and induces defence responses against green mould in citrus fruit. Postharvest Biology and Technology, 115, 18– 29.

Fan, J., Crooks, C. & Lamb, C. (2008) High-throughput quantitative luminescence assay of the growth in planta of *Pseudomonas syringae* chromosomally tagged with *Photorhabdus luminescens* luxCDABE. Plant Journal, 53, 393–399.

Fan, Y., Lu, X., Chen, X., Wang, J., Wang, D., Wang, S. et al. (2021) Cotton transcriptome analysis reveals novel biological pathways that eliminate reactive oxygen species (ROS) under sodium bicarbonate (NaHCO_3_) alkaline stress. Genomics, 113, 1157–1169.

Flora, G. & Rani, S.M.V. (2012) An approach towards control of blast by foliar application of seaweed concentrate. Science Research Reporter, 2, 213–217.

Ganiyu, S.A., Popoola, A.R., Owolade, O.F. & Fatona, K.A. (2017) Control of common bacterial blight disease of cowpea (*Vigna unguiculata* [L.] Walp) with certain plant extracts in Abeokuta, Nigeria. Journal of Crop Improvement, 31, 280–288.

German, L., Yeshvekar, R. & Benitez-Alfonso, Y. (2023) Callose metabolism and the regulation of cell walls and plasmodesmata during plant mutualistic and pathogenic interactions. Plant Cell & Environment, 46, 391–404.

Görlach, J., Volrath, S., Knauf-Beiter, G., Hengy, G., Beckhove, U., Kogel, K.H. et al. (1996) Benzothiadiazole, a novel class of inducers of systemic acquired resistance, activates gene expression and disease resistance in wheat. Plant Cell, 8, 629–643.

Habibullah, M., Sumardiyono, C. & Widiastuti, A. (2020) Potency of non-fungicide chemicals for maize inducing resistance against downy mildew. Jurnal Perlindungan Tanaman Indonesia, 24, 154–160.

Hael-Conrad, V., Perato, S.M., Arias, M.E., Martínez-Zamora, M.G., Di Peto, P.D.L.Á., Martos, G.G. et al. (2018) The elicitor protein AsES induces a systemic acquired resistance response accompanied by systemic microbursts and micro–hypersensitive responses in *Fragaria ananassa*. Molecular Plant-Microbe Interactions, 31, 46–60.

Hankins, S.D. & Hockey, H.P. (1990) The effect of a liquid seaweed extract from *Ascophyllum nodosum* (Fucales, Phaeophyta) on the two-spotted red spider mite *Tetranychus urticae*. In: Lindstrom, S.C. & Gabrielson, P.W. (Eds) Proceedings of the Thirteenth International Seaweed Symposium, 13–18 August 1989, Vancouver, BC: Springer, pp. 555–559.

Hawrylak-Nowak, B., Dresler, S., Stasińska-Jakubas, M., Wójciak, M., Sowa, I. & Matraszek-Gawron, R. (2021) NaCl-induced elicitation alters physiology and increases accumulation of phenolic compounds in *Melissa officinalis* L. International Journal of Molecular Sciences, 22, 6844.

Health Canada. (2025) Search Product Label. Pest Management Regulatory Agency, Health Canada, Ottawa, ON, Canada. Available from https://pr-rp.hc-sc.gc.ca/ls-re/index-eng.php. [Accessed 20th May 2025]

Hermann, S., Orlik, M., Boevink, P., Stein, E., Scherf, A., Kleeberg, I. et al. (2022) Biocontrol of plant diseases using *Glycyrrhiza glabra* leaf extract. Plant Disease, 106, 3133–3144.

Hervieux, V., Yaganza, E.S., Arul, J. & Tweddell, R.J. (2002) Effect of organic and inorganic salts on the development of *Helminthosporium solani*, the causal agent of potato silver scurf. Plant Disease, 86, 1014–1018.

Horst, R.K., Kawamoto, S.O. & Porter, L.L. (1992) Effect of sodium bicarbonate and oils on the control of powdery mildew and black spot of roses. Plant Disease, 76, 247–251.

Hussein, R.M. & Ahmed, F.A. (2022) Evaluation of the antibacterial and antibiofilm activities of sodium benzoate against the soft rot disease of tomato in Iraq. Archives of Phytopathology and Plant Protection, 55, 636–651.

Islam, M.T., Gan, H.M., Ziemann, M., Hussain, H.I., Arioli, T. & Cahill, D. (2020) Phaeophyceaean (brown algal) extracts activate plant defense systems in *Arabidopsis thaliana* challenged with *Phytophthora cinnamomi*. Frontiers in Plant Science, 11, 852.

Jacob, C., Panchal, S. & Melotto, M. (2017) Surface inoculation and quantification of *Pseudomonas syringae* population in the *Arabidopsis* leaf apoplast. Bio-Protocol, 7, e2167.

Jayaraman, J., Norrie, J. & Punja, Z.K. (2011) Commercial extract from the brown seaweed *Ascophyllum nodosum* reduces fungal diseases in greenhouse cucumber. Journal of Applied Phycology, 23, 353–361.

Jeandet, P., Adrian, M., Breuil, A.C., Debord, S., Sbaghi, M., Joubert, J.M. et al. (2000) Chemical induction of phytoalexin synthesis in grapevines: Application to the control of grey mould in the vineyard. Acta Horticulturae, 528, 591–596.

Jia, X., Zeng, H., Wang, W., Zhang, F. & Yin, H. (2018) Chitosan oligosaccharide induces resistance to *Pseudomonas syringae* pv. *tomato* DC3000 in *Arabidopsis thaliana* by activating both salicylic acid–and jasmonic acid–mediated pathways. Molecular Plant-Microbe Interactions, 31, 1271–1279.

Karabulut, O.A., Smilanick, J.L., Gabler, F.M., Mansour, M. & Droby S. (2003) Near-harvest applications of *Metschnikowia fructicola*, ethanol, and sodium bicarbonate to control postharvest diseases of grape in central California. Plant Disease, 87, 1384–1389.

Kesawat, M.S., Satheesh, N., Kherawat, B.S., Kumar, A., Kim, H.U., Chung, S.M. et al. (2023) Regulation of reactive oxygen species during salt stress in plants and their crosstalk with other signaling molecules–Current perspectives and future directions. Plants, 12, 864.

Kishore, G.K., Pande, S. & Harsha, S. (2007) Evaluation of essential oils and their components for broad-spectrum antifungal activity and control of late leaf spot and crown rot diseases in peanut. Plant Disease, 91, 375–379.

Kohler, A., Schwindling, S. & Conrath, U. (2002) Benzothiadiazole-induced priming for potentiated responses to pathogen infection, wounding, and infiltration of water into leaves requires the *NPR1/NIM1* gene in Arabidopsis. Plant Physiology, 128, 1046–1056.

Konstantinidou-Doltsinis, S., Markellou, E., Kasselaki, A.M., Fanouraki, M.N., Koumaki, C.M., Schmitt, A. et al. (2006) Efficacy of Milsana®, a formulated plant extract from *Reynoutria sachalinensis*, against powdery mildew of tomato (*Leveillula taurica*). Biocontrol, 51, 375– 392.

Koornneef, A., Verhage, A., Leon-Reyes, A., Reinier, S., Van Loon, L.C. & Pieterse, C.M.J. (2008) Towards a reporter system to identify regulators of cross-talk between salicylate and jasmonate signaling pathways in Arabidopsis. Plant Signaling & Behavior, 3, 543–546.

Lefevere, H., Bauters, L. & Gheysen, G. (2020) Salicylic acid biosynthesis in plants. Frontiers in Plant Science, 11, 338.

Liu, N.J., Zhang, T., Liu, Z.H., Chen, X., Guo, H.S., Ju, B.H. et al. (2020) Phytosphinganine affects plasmodesmata permeability via facilitating PDLP5-stimulated callose accumulation in *Arabidopsis*. Molecular Plant, 13, 128–143.

Llorens, E., Mateu, M., González-Hernández, A.I., Agustí-Brisach, C., García-Agustín, P., Lapeña, L. et al. (2019) Extract of *Mimosa tenuiflora* and Quercus robur as potential eco-friendly management tool against *Sclerotinia sclerotiorum* in *Lactuca sativa* enhancing the natural plant defences. European Journal of Plant Pathology, 153, 1105–1118.

Lyousfi, N., Letrib, C., Legrifi, I., Blenzar, A., El Khetabi, A., El Hamss, H. et al. (2022) Combination of sodium bicarbonate (SBC) with bacterial antagonists for the control of brown rot disease of fruit. Journal of Fungi, 8, 636.

Mahizan, N.A., Yang, S.-K., Moo, C.-L., Song, A.A.-L., Chong, C.M., Chong, C.-W. et al. (2019) Terpene derivatives as a potential agent against antimicrobial resistance (AMR) pathogens. Molecules, 24, 2631.

Mahmood, B., Tariq-Khan, M., Azad, A., Rahim, N., Arif, S. & Khan, M.R. et al. (2022) Management of late blight of potato caused by *Phytophthora infestans* through botanical aqueous extracts. International Journal of Phytopathology, 11, 35–43.

Margaritopoulou, T., Toufexi, E., Kizis, D., Balayiannis, G., Anagnostopoulos, C., Theocharis, A. et al. (2020) *Reynoutria sachalinensis* extract elicits SA-dependent defense responses in courgette genotypes against powdery mildew caused by *Podosphaera xanthii*. Scientific Reports, 10, 3354.

Martins, P.M.M., Merfa, M.V., Takita, M.A. & De Souza, A.A. (2018) Persistence in phytopathogenic bacteria: Do we know enough? Fronbers in Microbiology, 9, 1099.

Masami, N., Masao, G., Katsumi, A. & Tadaaki, H. (2004) Nucleotide sequence and organization of copper resistance genes from *Pseudomonas syringae* pv. *acbnidiae*. European Journal of Plant Pathology, 110, 223–226.

Mecteau, M.R., Arul, J. & Tweddell, R.J. (2002) Effect of organic and inorganic salts on the growth and development of *Fusarium sambucinum*, a causal agent of potato dry rot. Mycological Research, 106, 688–696.

Mesquita, A.F., Gonçalves, F.J.M. & Gonçalves, A.M.M. (2023) The lethal and sub-lethal effects of fluorinated and copper-based pesticides—A review. International Journal of Environmental Research and Public Health, 20, 3706.

Mimouni, S., Delisle-Houde, M., Demers, F., Filion, M. & Tweddell, R.J. (2025) Maple leaf extracts to control angular leaf spot of cucurbits caused by *Pseudomonas syringae*. Plant Pathology, 74, 1114–1120.

Mirik, M., Aysan, Y. & Cinar, O. (2007) Copper-resistant strains of *Xanthomonas axanopodis* pv. *vesicatoria* (Doidge) dye in the eastern Mediterranean region of Turkey. Journal of Plant Pathology, 89, 153–154.

Montesinos-Herrero, C., Moscoso-Ramírez, P.A. & Palou, L. (2016) Evaluation of sodium benzoate and other food additives for the control of citrus postharvest green and blue molds. Postharvest Biology and Technology, 115, 72–80.

Moschetto, F.A., Lopes, M.F., Silva, B.P. & Neto, M.C.L. (2019) Sodium benzoate inhibits germination, establishment, and development of rice plants. Theoretical and Experimental Plant Physiology, 31, 377–385.

Nashwa, S.M.A. & Abo-Elyousr, K.A.M. (2012) Evaluation of various plant extracts against the early blight disease of tomato plants under greenhouse and field conditions. Plant Protection Science, 48, 74–79.

Naz, R., Bano, A., Nosheen, A., Yasmin, H., Keyani, R., Shah, S.T.A. et al. (2021) Induction of defense-related enzymes and enhanced disease resistance in maize against *Fusarium verticillioides* by seed treatment with *Jacaranda mimosifolia* formulations. Scientific Reports, 11, 59.

Nehela, Y., Taha, N.A., Elzaawely, A.A., Xuan, T.D., Amin, M.A., Ahmed, M.E. et al. (2021) Benzoic acid and its hydroxylated derivatives suppress early blight of tomato (*Alternaria solani*) via the induction of salicylic acid biosynthesis and enzymatic and nonenzymatic antioxidant defense machinery. Journal of Fungi, 7, 663.

Nguefack, J., Nguikwie, S.K., Fotio, D., Dongmo, B., Zollo, P.H.A., Leth, V. et al. (2007) Fungicidal potential of essential oils and fractions from *Cymbopogon citratus, Ocimum gratissimum* and *Thymus vulgaris* to control *Alternaria padwickii* and *Bipolaris oryzae*, two seed-borne fungi of rice (*Oryza Sativa* L). Journal of Essential Oil Research, 19, 581–587.

Olivier, C., MacNeil, C.R. & Loria, R. (1999) Application of organic and inorganic salts to field grown potato tubers can suppress silver scurf during potato storage. Plant Disease, 83, 814– 818.

Olivier, C., Halseth, D.E., Mizubuti, E.S.G. & Loria, R. (1998) Postharvest application of organic and inorganic salts for suppression of silver scurf on potato tubers. Plant Disease, 82, 213–217.

Palmer, C.L., Horst, R.K. & Langhans, R.W. (1997) Use of bicarbonates to inhibit in vitro colony growth of *Botrytis cinerea*. Plant Disease, 81, 1432–1438.

Palou, L., Smilanick, J.L. & Crisosto, C.H. (2009) Evaluation of food additives as alternative or complementary chemicals to conventional fungicides for the control of major postharvest diseases of stone fruit. Journal of Food Protection, 72, 1037–1046.

Peng, Y., Yang, J., Li, X. & Zhang, Y. (2021) Salicylic acid: biosynthesis and signaling. Annual Review of Plant Biology, 72, 761–791.

Pradhanang, P.M., Momol, M.T., Olson, S.M. & Jones, J.B. (2003) Effects of plant essential oils on *Ralstonia solanacearum* population density and bacterial wilt incidence in tomato. Plant Disease, 87, 423–427.

Punja, Z.K. & Gaye, M.M. (1993) Influence of postharvest handling practices and dip treatments on development of black root rot on fresh market carrots. Plant Disease, 77, 989–995.

Qin, P., Wei, A., Zhao, D., Yao, Y., Yang, X., Dun, B. et al. (2017) Low concentration of sodium bicarbonate improves the bioactive compound levels and antioxidant and a-glucosidase inhibitory activities of tartary backwheat sprouts. Food Chemistry, 224, 124–130.

Ragab, M.M., Ashour, A.M.A., Abdel-Kader, M.M., El-Mohamady, R. & Abdel-Aziz, A. (2012) *In vitro* evaluation of some fungicides alternatives against *Fusarium oxysporum* the causal of wilt disease of pepper (*Capsicum annum* L.). International Journal of Agriculture and Forestry, 2, 70–77.

Rengel, Z. (1992) Role of calcium in aluminum toxicity. New Phytologist, 121, 499–513.

Renick, L.J., Cogal, A.G. & Sundin, G.W. (2008) Phenotypic and genetic analysis of epiphytic *Pseudomonas syringae* populations from sweet cherry in Michigan. Plant Disease, 92, 372– 378.

Rishad, M.B., Sultana, A., Chakraborty, S., Humaira, R.K. & Khokon, M.A.R. (2021) Effect of foliar application of salicylic acid, chitosan and benzoic acid in elevating total phenol and H_2_O_2_ content in rice leaves to modulate resistance against blast disease. Bangladesh Journal of Plant Pathology, 37, 7–14.

Schoffer, J.T., Sauvé, S., Neaman, A. & Ginocchio, R. (2020) Role of leaf litter on the incorporation of copper-containing pesticides into soils under fruit production: a review. Journal of Soil Science and Plant Nutrition, 20, 990–1000.

Seo, P.J., Lee, A.-K., Xiang, F. & Park, C.-M. (2008) Molecular and functional profiling of *Arabidopsis pathogenesis-related* genes: Insights into their roles in salt response of seed germination. Plant Cell & Physiology, 49, 334–344.

Seo, S.Y., Wi, S.J. & Park, K.Y. (2020) Functional switching of NPR1 between chloroplast and nucleus for adaptive response to salt stress. Scientific Reports, 10, 4339.

Shafique, S., Shafique, S., Zameer, M. & Asif, M. (2019) Plant defense system activated in chili plants by using extracts from *Eucalyptus citriodora*. Biocontrol Science, 24, 137–144.

Shaker, A.S., Marrez, D.A., Ali, M.A. & Fathy, H.M. (2022) Potential synergistic effect of *Alhagi graecorum* ethanolic extract with two conventional food preservatives against some foodborne pathogens. Archives of Microbiology, 204, 686.

Sholberg, P.L., Bedford, K.E., Haag, P. & Randall, P. (2001) Survey of *Erwinia amylovora* isolates from British Columbia for resistance to bactericides and virulence on apple. Canadian Journal of Plant Pathology, 23, 60–67.

Sivakumar, D., Hewarathgamagae, N.K., Wijeratnam, R.S.W. & Wijesundera, R.L.C. (2002) Effect of ammonium carbonate and sodium bicarbonate on anthracnose of papaya. Phytoparasitica, 30, 486–492.

Smilanick, J.L., Margosan, D.A., Mlikota, F., Usall, J. & Michael, I.F. (1999) Control of citrus green mold by carbonate and bicarbonate salts and the influence of commercial postharvest practices on their efficacy. Plant Disease, 83, 139–145.

Soliman, A.S., Ahmed, A.Y., Abdel-Ghafour, S.E., El-Sheekh, M.M. & Sobhy, H.M. (2018) Antifungal bio-efficacy of the red algae *Gracilaria confervoides* extracts against three pathogenic fungi of cucumber plant. Middle East Journal of Applied Science, 8, 727–735.

Soltaniband, V., Barrada, A., Delisle-Houde, M., Dorais, M., Tweddell, R.J. & Michaud, D. (2024) Forest tree extracts induce resistance to *Pseudomonas syringae* pv. *tomato* in Arabidopsis. Scientific Reports, 14, 24726.

Stanojević, D., Comic, L.J., Stefanovic, O. & Sukdolak, S.S. (2010) *In vitro* synergistic antibacterial activity of *Melissa officinalis* L. and some preservatives. Spanish Journal of Agricultural Research, 8, 109–115.

Subramanian, S., Sangha, J.S., Gray, B.A., Singh, R.P., Hiltz, D., Critchley, A.T. et al. (2011) Extracts of the marine brown macroalga, *Ascophyllum nodosum*, induce jasmonic acid dependent systemic resistance in *Arabidopsis thaliana* against *Pseudomonas syringae* pv. *tomato* DC3000 and *Sclerotinia sclerotiorum*. European Journal of Plant Pathology, 131, 237–248.

Thomma, B.P., Eggermont, K., Penninckx, I.A., Mauch-Mani, B., Vogelsang, R., Cammue, B.P. et al. (1998) Separate jasmonate-dependent and salicylate-dependent defense-response pathways in *Arabidopsis* are essential for resistance to distinct microbial pathogens. Proceedings of the National Academy of Sciences U.S.A., 95, 15107–15111.

Tiku, A.R. (2020) Antimicrobial compounds (phytoanticipins and phytoalexins) and their role in plant defense. In: Mérillon, J.M. & Ramawat, K. (Eds.) Co-Evolution of Secondary Metabolites. Cham, Switzerland: Springer, pp. 845–868.

Tremblay, V., Delisle-Houde, M., Demers, F., D’Amours, C., Filion, M. & Tweddell, R.J. (2024) Sugar maple leaf extracts: A new tool to control bacterial canker of tomato caused by *Clavibacter michiganensis* subsp. *michiganensis*. Plant Pathology, 73, 2123–2131.

Ullah, C., Chen, Y.-H., Ortega, M.A. & Tsai, C.-J. (2023) The diversity of salicylic acid biosynthesis and defense signaling in plants: Knowledge gaps and future opportunities. Current Opinion in Plant Biology, 72, 102349.

Vandesompele, J., De Preter, K., Pattyn, F., Poppe, B., Van Roy, N., De Paepe, A. et al. (2002) Accurate normalization of real-time quantitative RT-PCR data by geometric averaging of multiple internal control genes. Genome Biology, 3, research0034.1.

Vanneste, J.L., McLaren, G.F., Yu, J., Cornish, D.A. & Boyd, R. (2005) Copper and streptomycin resistance in bacterial strains isolated from stone fruit orchards in New Zealand. New Zealand Plant Protection, 58, 101–105.

Vilaplana, R., Alba, P. & Valencia-Chamorro, S. (2018) Sodium bicarbonate salts for the control of postharvest black rot disease in yellow pitahaya (*Selenicereus megalanthus*). Crop protection, 114, 90–96.

Wang, X., Sager, R., Cui, W., Zhang, C., Lu, H. & Lee, J.Y. (2013) Salicylic acid regulates plasmodesmata closure during innate immune responses in *Arabidopsis*. Plant Cell, 25, 2315–2329.

Williams, M., Senaratna, T., Dixon, K. & Sivasithamparam, K. (2003) Benzoic acid induces tolerance to biotic stress caused by *Phytophthora cinnamomi* in *Banksia attenuata*. Plant Growth Regulation, 41, 89–91.

Xin, X.-F. & He, S.Y. (2013) *Pseudomonas syringae* pv. *tomato* DC3000: a model pathogen for probing disease susceptibility and hormone signaling in plants. Annual Review of Phytopathology, 51, 473–498.

Xin, X.-F., Kvitko, B. & He, S.Y. (2018) *Pseudomonas syringae*: what it takes to be a pathogen. Nature Reviews in Microbiology, 16, 316–328.

Yaganza, E.S., Tweddell, R.J. & Arul, J. (2009) Physicochemical basis for the inhibitory effects of organic and inorganic salts on the growth of *Pectobacterium carotovorum* subsp. *carotovorum* and *Pectobacterium atrosepticum*. Applied and Environmental Microbiology, 75, 1465–1469.

Yaganza, E.S., Tweddell, R.J. & Arul, J. (2014) Postharvest application of organic and inorganic salts to control potato (*Solanum tuberosum* L.) storage soft rot: plant tissue–salt physicochemical interactions. Journal of Agricultural and Food Chemistry, 62, 9223–9231.

Youssef, K., Roberto, S.R., Tiepo, A.N., Constantino, L.V., de Resende, J.T.V. & Abo-Elyousr, K.A.M. (2020) Salt solution treatments trigger antioxidant defense response against gray mold disease in table grapes. Journal of Fungi, 6, 179.

Youssef, K., Sanzani, S.M., Ligorio, A., Ippolito, A. & Terry, L.A. (2014) Sodium carbonate and bicarbonate treatments induce resistance to postharvest green mould on citrus fruit. Postharvest Biology and Technology, 87, 61–69.

Zhao, S. & Fernald, R.D. (2005) Comprehensive algorithm for quantitative real-time polymerase chain reaction. Journal of Computational Biology, 12, 1047–1064.

Zhou, H., Shi, H., Yang, Y., Feng, X., Chen, X., Xiao, F. et al. (2024) Insights into plant salt stress signaling and tolerance. Journal of Genetics and Genomics, 51, 16–34.

Zimmermann, P., Hirsch-Hoffmann, M., Hennig, L. & Gruissem, W. (2004) GENEVESTIGATOR. Arabidopsis microarray database and analysis toolbox. Plant Physiology, 136, 2621–2632.

Ziv, O. & Zitter, T.A. (1992) Effects of bicarbonates and film-forming polymers on cucurbit foliar diseases. Plant Disease, 76, 513–517.

Zuo, G.Y., Han, Z.Q., Hao, X.Y., Han, J., Li, Z.S. & Wang, G.C. (2014) Synergy of aminoglycoside antibiotics by 3-Benzylchroman derivatives from the Chinese drug *Caesalpinia sappan* against clinical methicillin-resistant *Staphylococcus aureus* (MRSA). Phytomedicine, 21, 936– 941.

Zuzarte, M., Vale-Silva, L., Gonçalves, M.J., Cavaleiro, C., Vaz, S., Canhoto, J. et al. (2012) Antifungal activity of phenolic-rich *Lavandula multifida* L. essential oil. European Journal of Clinical Microbiology & Infectious Diseases, 31, 1359–1366.

