## Supplementary Figures for "GRAS salts and eastern hemlock extract: a dual approach to sustainable plant disease management"

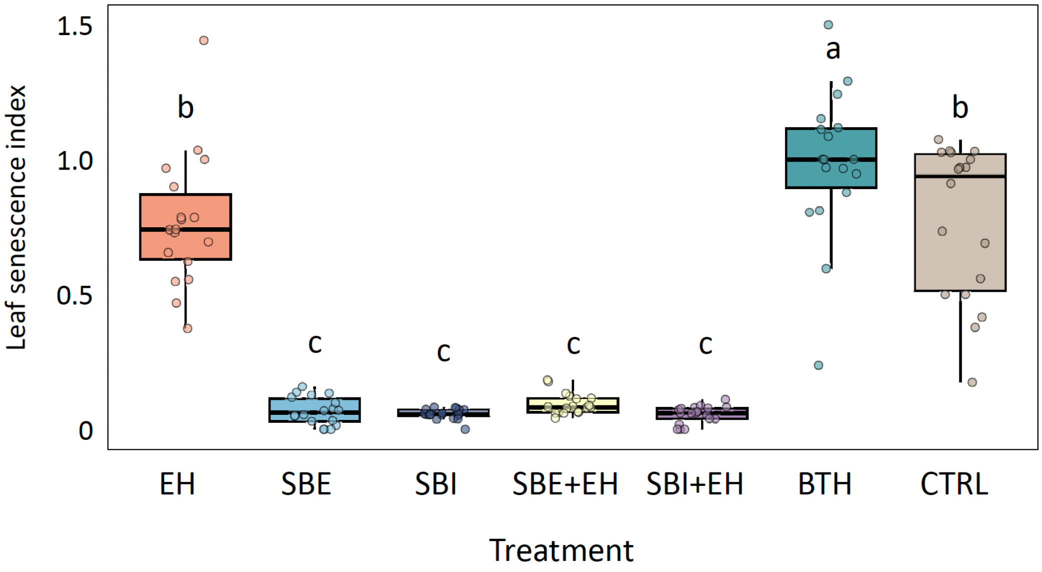


**Supplementary Figure S1.** Leaf senescence index of Arabidopsis plants treated with sodium benzoate (SBE) or sodium bicarbonate (SBI), alone or combined with the eastern hemlock (EH) extract. The plants were treated with 0.01 M SBE or 0.01 M SBI, with the EH extract at 25 mg/mL, or with SBE or SBI in combination with the EH extract at the same working concentrations. Sterile water was used as a negative control treatment (CTRL), ACTIGARD$^{TM}$ 50WG (BTH) at 0.5 mg/mL as a non-toxic positive control for SA signaling induction. Treatments were applied by foliar spraying and the leaf samples collected after nine days, with no bacterial inoculation two days after spraying. Individual measurements are shown as dots. In each box, the horizontal line shows the median, the lower and upper edges the first (Q1) and third (Q3) quartiles, respectively. Treatments sharing the same letter are not significantly different (n=18; *post-hoc* pairwise multiple comparisons of estimated marginal means with Tukey's adjustment, with an alpha threshold value of 5%).

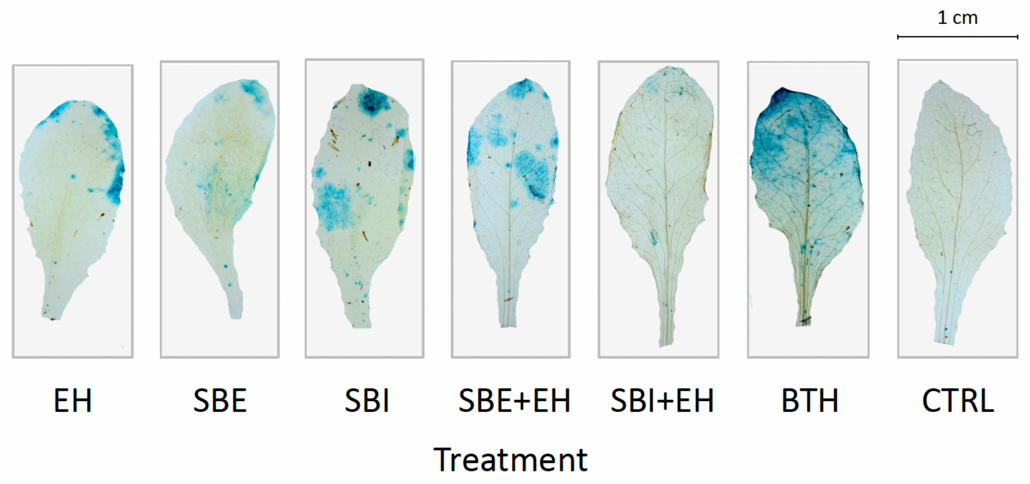


**Supplementary Figure S2.** Representative GUS staining patterns in leaves of Arabidopsis reporter line *PR1::GUS* treated with sodium benzoate (SBE) or sodium bicarbonate (SBI), alone or combined with the eastern hemlock (EH) extract. The plants were treated with 0.01 M SBE or 0.01 M SBI, with the EH extract at 25 mg/mL, or with SBE or SBI combined with the EH extract at the same working concentrations. Sterile water was used as a negative control treatment (CTRL), ACTIGARD$^{TM}$ 50WG (BTH) at 0.5 mg/mL as a non-toxic positive control for SA signaling induction. Treatments were applied by foliar spraying two days prior to inoculation with *Pst* DC3000, and the leaf samples collected after nine days.
