## Supplementary Table S1 for "GRAS salts and eastern hemlock extract: a dual approach to sustainable plant disease management"

**Supplementary Table S1.** DNA primers for the RT-qPCR assay.

| Target gene | Locus number [1] | | Primer sequence (5′ → 3′) |
|---|---|---|---|
| *AtPR1* | at2g14610 | F | TTCTTCCCTCGAAAGCTCAA |
| | | R | AAGGCCCACCAGAGTGTATG |
| *AtLOX2* | at3g45140 | F | ATTACGGTAGAAGACTACGCACAAC |
| | | R | GTAATTTAAGCTCTACCCCCTTGAG |
| *AtPR3* | at3g12500 | F | GGCCAGACTTCCCATGAAAC |
| | | R | CTTGAAACAGTAGCCCCATGAA |

[1] Accession numbers from The Arabidopsis Information Resource database (www.arabidopsis.org).
